# Metagenomic polymorphic toxin effector and immunity profiling predicts microbiome development and disease-related dysbiosis

**DOI:** 10.1101/2025.07.08.662037

**Authors:** Hunter W Schroer, Francesco Beghini, Juan Antonio Raygoza Garay, Nicholas A Christakis, Dustin E. Bosch

## Abstract

Bacteria use antagonistic interbacterial weapons such as polymorphic toxin secretion systems (TSS) to compete for niches in the human gut microbiome. We developed a bioinformatic marker gene approach (PolyProf) to quantify TSS including ∼200 effector and immunity genes and applied it to ∼15,000 publicly available human metagenomes. PolyProf alpha and beta diversity readily distinguished 12 different human disease states. Decision tree machine learning models integrating bacterial taxonomy with PolyProf had near-perfect accuracy (ROC area 1.00) for all 12 disease states. During microbiome development in the first year of life, PolyProf alpha diversity increases, and beta diversity becomes increasingly like the maternal microbiome, influenced by vertical transfer, delivery mode, and breastfeeding. PolyProf is related to strain sharing among adults through social interactions. In summary, interbacterial antagonism with TSS shapes microbiome development and interpersonal strain sharing. Since PolyProf distinguishes diverse adult disease statuses, these dynamics may contribute to non-genetic inheritance.

## Introduction

Bacteria use a complex arsenal of toxins to compete for niches in ecosystems with limited resources ^2,3^. From a bacterial perspective, the human gut microbiome is a densely populated ecosystem with environmental pressures such as changing dietary nutrition sources and host factors like innate and adaptive immunity and secretion of antimicrobial peptides ^4^. While intestinal bacteria have commensal relationships with the human host, interbacterial relationships are more complex, ranging from competition/exploitation to mutualism ^5^. The complex interbacterial interactions can in turn impact the host, and the gut microbiome has a well-established bi-directional relationship with host disease.

For example, in inflammatory bowel disease (IBD), acute and chronic intestinal inflammation, epithelial barrier compromise, and altered motility of the fecal stream all significantly alter the intestinal ecosystem, which naturally results in changes to bacterial communities away from the “healthy” state, referred to as dysbiosis ^6^. Although dysbiosis is often an effect of disease, causal links for the dysbiotic microbiome in the pathogenesis and progression of several diseases has also been established. For example, germ-free conditions prevent the development of colitis in mouse models of IBD, and the transfer of bacteria from diseased animals or IBD patients exacerbates colitis ^6^. Similarly, fecal transplants from obese humans are sufficient to promote weight gain in mice ^7^. Mechanisms include bacterial community modulation of host immunity and production of disease-promoting proteins and metabolites ^8–10^.

Development of the gut microbiome in infancy is a highly dynamic process that involves the acquisition of bacteria from the environment and rapidly changing host factors such as immune system development ^11–14^. A recurring theme in infant microbiome studies is a predominance of vertical transfer from close-contact caretakers. In the case of maternal primary caregiving and/or breastfeeding, there is a tendency of the infant gut microbiome to become increasingly like the maternal gut microbiome ^11,14^. Vertical strain transfer has been demonstrated in several studies to occur at birth and throughout the first year of life through nutritional sources and close contact ^15–19^. In addition, infant microbiome development is influenced rather dramatically by delivery mode (vaginal delivery [SVD] *vs.* Cesarean section [CSD]). Recent trials have demonstrated that intentional exposure of infants to maternal commensal bacteria can accelerate the convergence of the infant and maternal microbiomes, highlighting the feasibility of intervention in the developing microbiome ^20^. However, the long-term health benefits of these interventions remain to be established. Since dysbiosis is a prominent feature in many diseases, and dysbiotic microbiomes play causal roles in several, it is distinctly possible that vertical transmission of gut microbiome components may underly non-genetic disease inheritance patterns ^21,22^.

Polymorphic toxin effectors utilized in interbacterial antagonism compromise central processes required for cell survival, such as the maintenance of membrane, cell wall, and genome integrity ^2^. Most effectors are highly efficient enzymes, such that delivery of one or a few effector molecules is fatal to a non-immune recipient bacterium. In the interbacterial antagonism arms race, polymorphic effector genes have proliferated. Because of high sequence diversity, frequent horizontal transfer, and technical challenges in molecular studies of toxic proteins, bioinformatic identification and function prediction of polymorphic toxin effectors is difficult ^3,23,24^.

Our approach to profiling here relies on the Zhang et al. comprehensive bioinformatic analysis that identified at least 150 distinct effector domains with highly diverse predicted catalytic activities ^3^. Effector-encoding cells protect themselves and kin from intoxication with immunity proteins, often encoded immediately 3’ on the genome from the effector gene. Immunity can be acquired by inheritance or horizontal transfer of toxin secretion systems, many of which are transferrable via plasmids or other mobile genetic elements ^25–27^. Another defensive strategy is the acquisition of “orphan” immunity genes without the corresponding effector or energetically costly secretion system ^25,28^. For example, Bacteroidales use genetically mobile arrays of immunity genes or acquired interbacterial defense systems (AID) to confer protection from several effector classes ^28^.

Polymorphic toxin effectors are delivered to recipient competitors through several conserved mechanisms ^2^. For example, colicins are synthesized and released from dying *E. coli* and co-opt membrane transporters to enter and recipient bacteria ^2^. The contact-dependent growth inhibition (CDI) or type V secretion system (T5SS) delivers effectors to contacting cells through cell surface receptor interactions ^2^. The type VI secretion system (T6SS) forms an apparatus structurally related to bacteriophage, comprised of TssA-R proteins ^2,29^. A contractile sheath propels a tip structure decorated with an effector(s) to deliver effectors into the recipient periplasm and/or cytoplasm ^30^. T6SS are divided into genetically related clades with different taxonomic distributions, and T6SS^iii^ is exclusively distributed among Bacteroidales ^31^. T6SS^iii^ membrane complexes are composed of TssQ and other structural proteins that are lacking in the T6SS found in Proteobacteria ^32^. Esx or type VII secretion systems (ESS / T7SS) were initially described in Mycobacteria, and homologs are widespread in Bacillota ^33^. EssB is one of several conserved integral membrane proteins required for secretion ^2^.

Polymorphic toxin secretion systems have predominantly been discovered and initially studied in pathogens and environmental bacteria. However, T6SS^iii^ are known to mediate interbacterial antagonism and selective colonization in the human gut microbiome ^28,31,34^. Inactivation of T6SS in Bacteroidales impairs competitive growth *in vitro* and competitive colonization of gnotobiotic mice ^28,31,34^. A human gut metagenome analysis of T6SS^iii^ and associated effector/immunity genes demonstrated the GA3 architecture systems found in *Bacteroides fragilis* are associated with increased *Bacteroides* abundance and are enriched in infant microbiomes ^35^. Dominant effector types are proposed to competitively exclude colonization by related, but non-immune Bacteroidales ^27,28,36,37^, potentially contributing to individual microbiome stability. Much less is known regarding the roles of toxin secretion systems outside of Bacteroidales in the gut microbiome context. The environmental pressures driving disease-related dysbiosis likely enhance the utility of competitive antagonism. However, the relationship between polymorphic toxin secretion systems in the gut microbiome and human diseases remains an open area of inquiry. For instance, our recent study of a nuclease effector Tde (encoded with T6SS^iii^ in Bacteroidales isolated from humans) found that effector and T6SS^iii^ gene abundances in metagenomes were associated with IBD diagnoses ^25^.

The prior research provides proof-of-concept that polymorphic toxin secretion genes may be disease-specific markers of dysbiosis. The model of competitive exclusion of non-immune Bacteroidales strains ^35^ and the prominence of this order in the effects of delivery mode on infant microbiomes also suggests that polymorphic toxins may be an important component of gut microbiome development ^12,14,35^. We hypothesized that polymorphic toxin secretion system genes play a role in infant microbiome development and are a feature of disease-related dysbiosis. Therefore, the primary goals of this study are to 1) develop a bioinformatics pipeline (PolyProf) for quantitation of polymorphic toxin secretion genes in metagenomes, 2) describe the landscape of PolyProf in health, development, and disease, 3) test the predictive value of PolyProf for human disease diagnosis, and 4) identify highly disease-associated polymorphic toxin genes for future mechanistic studies.

## Results

### Development of a metagenomic polymorphic toxin effector/immunity profiling pipeline (PolyProf)

To identify disease associations with intestinal microbiome effectors and immunity genes, we constructed a marker protein sequence database for use with HUMAnN ^38^. We restricted the initial database to the effector and immunity classes described by Zhang et al. ^3^ (Table S1, Figure S1). Inclusion of the T6SS^iii^ structural proteins allowed validation testing by co-abundance, because all 13 targets are required to assemble a functional apparatus in any Bacteroidales. High positive correlation of T6SS^iii^ marker abundances were observed across all datasets (Figure S2), reflecting the expected co-abundance. Effector immunity pairs are also expected to co-occur in metagenomes, as observed for Tde/Tdi (Figure S2). However, strong linear effector/immunity co-abundance is not expected due to the preponderance of orphan immunity in the gut microbiome ^28^. Conversely, co-abundance of markers not related to direct effector/immunity interactions may arise, for example, due to genetic mobility of secretion systems and immunity gene arrays (AIDs) ^27,28,36^.

PolyProf database marker performance was assessed using simulated metagenomic data ^39^, generated from randomly selected MAGs with and without the corresponding effector/immunity genes (Figure S3). Most PolyProf marker abundance measurements correlated linearly with relative abundance of the encoding MAGs. Exceptions were Ntox11 markers, which tended to underestimate (low sensitivity) true gene abundance, and TssR, Ntox30, LDpeptidase, and ToxREase4, which systematically overestimated abundance (low specificity).

We considered that PolyProf is likely influenced by taxonomy, because some effector/immunity and secretion system genes are restricted to specific taxonomic groups. To quantify and visualize taxonomic distributions of each marker using gut MAGs (MGnify), we generated trees and calculated enrichment Z-scores by taxonomic group (available online at PolyProf GitHub site). We then used 1537 metagenomes from the IBD cohorts (including many healthy controls) to calculate both PolyProf and taxonomic relative abundances using MetaPhlAn ^38^, followed by Bray-Curtis dissimilarity beta diversity NMDS analysis. The contribution of taxonomy to PolyProf diversity was assessed by plotting values from the dominant NMDS of each analysis and linear regression (Figure S4). An R^2^ value of 0.11 can be interpreted as approximately 11% of the variance in PolyProf being explained by taxonomy. To identify specific markers that are highly related to taxonomy, we predicted marker abundances from taxonomic relative abundances and the frequency of marker detection in MAGs derived from each family. The predicted abundances were compared to PolyProf measured abundances with linear regression to assess how much of the PolyProf marker abundance is explained by taxonomy (plots available online at PolyProf GitHub site). Ntox4 was most highly related to taxonomy (R^2^ = 0.42) in its restriction to Bacteroidales and Oscillospirales (Figure S4). The next eight markers most highly predictable from taxonomy were components of the Bacteroidales-specific T6SS^iii^ apparatus (R^2^ 0.20 – 0.24). The findings indicate that a small but significant fraction (∼11%) of PolyProf variance is due to differential abundance of bacterial taxa that tend to encode specific effector/immunity and secretion system genes. The influence of taxonomy on PolyProf is driven by a subset of the markers. For example, exclusion of the Bacteroidales-restricted T6SS^iii^ gene markers decreases the taxonomy-explained PolyProf variance from ∼11% to ∼7%. In biological terms, the low correlation between taxonomy and PolyProf reflects other contributions to effector/immunity abundances, such as selection for advantageous antagonism systems and horizontal transfer among diverse taxa. Because PolyProf and taxonomic beta diversity are not strongly correlated, they may have different and complementary predictive value in disease.

### Meta-analysis of large shotgun metagenomic gut microbiome datasets

We sought to identify effector/immunity diversity and abundance patterns in gut microbiomes that are associated with human disease. We applied PolyProf to ∼15,000 human fecal metagenomes grouped by one of 16 diagnoses (Figure S1, Table S2). A 3-fold analysis was then undertaken: 1) by diagnosis within a study, 2) each diagnosis compared to pooled adult “healthy controls” across studies, and 3) each diagnosis compared to all other metagenomes. The rationale for this approach was targeting effector/immunity profiles that are generalizable to populations with diverse health statuses. Global diversity of effector/immunity profiles exhibited significant differences by diagnosis (Figure 1). Some individual markers, including the DNAse effector/immunity pair Tde/Tdi ^25^ were independently predictive of disease state using multivariate logistic regression (Figure 1A). In prior research, *Phocaeicola vulgatus* was demonstrated to secrete Tde with a Bacteroidales-restricted T6SS^iii^ secretion system, conferring a competitive advantage over non-immune Bacteroidales derived from the same individual’s microbiome ^25^. To determine how taxonomy may influence PolyProf disease associations, we applied multivariable linear modeling (MaAsLin), with and without adjustment for Bacteroidales abundance (Figure S4). The relatively low T6SS^iii^ markers in infant microbiomes are influenced by Bacteroidales abundance, but their associations remain significant after adjustment for Bacteroidales. Other T6SS^iii^ disease associations, such as depletion in obesity and enrichment in ulcerative colitis, are not strongly affected by taxonomy (Figure S4).

**Figure 1.**
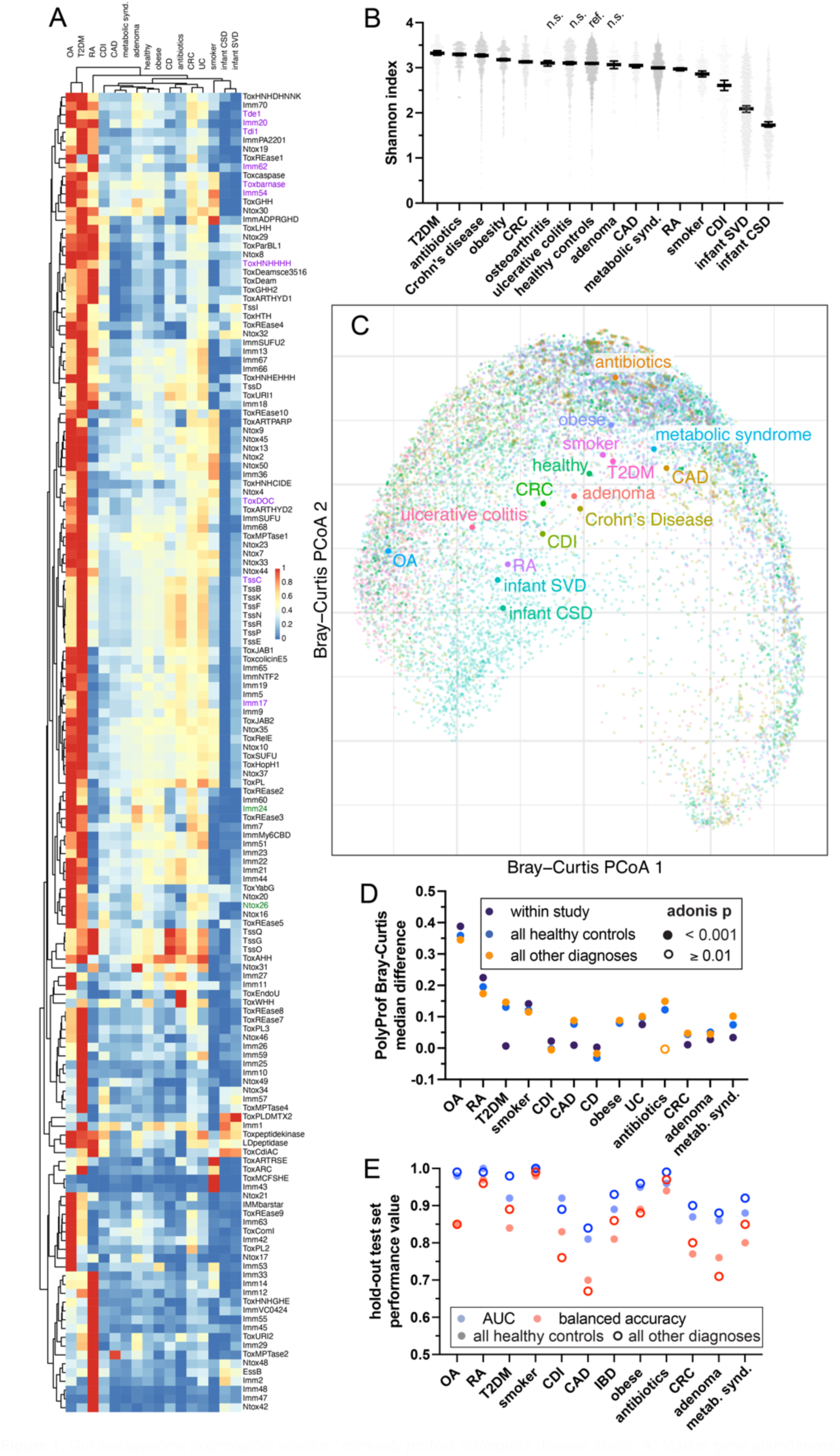
Gut metagenome polymorphic effector/immunity profiles distinguish disease states. A) Marker gene abundances, centered log-ratio transformed, are represented in a heat map with clustering by similarity. Osteoarthritis and type 2 diabetes (T2DM) subjects have highest overall abundance of many effector/immunity classes and T6SS^iii^. Independently diagnosis-distinguishing markers identified by logistic regression are colored green for p<0.001 and purple for p<0.01. B) Marker gene alpha diversity differs from “healthy control” comparators in all disease classes (adjusted t-test p<0.01) other than colon adenoma, ulcerative colitis, and osteoarthritis. C) Beta diversity also differed by diagnosis measured as Bray-Curtis dissimilarity PCoA. D) Beta diversity differences held (adonis2 p < 0.001) in a 3-tier analysis except for all diagnoses except the antibiotics-treated group. E) Decision tree machine learning (XGBoost) models were constructed to predict disease based on PolyProf. Performance is measured on a 20% hold-out test set with AUC and balanced accuracy. Individual ROC plots and top feature rankings for each diagnosis are in Figure S7-9. Abbreviations: OA osteoarthritis, RA rheumatoid arthritis, CSD Cesarean delivery, SVD vaginal delivery, CRC colorectal carcinoma, CD Crohn’s disease, UC ulcerative colitis, CDI *C. difficile* infection, met synd. metabolic syndrome.

Marker gene alpha diversity, measured as the Shannon index was either higher or lower than the “healthy controls” referent in all diagnostic groups except colorectal adenoma, ulcerative colitis (UC), and osteoarthritis (OA) (Figure 1B). The highest PolyProf alpha diversity is observed in type 2 diabetes (T2DM) metagenomes, which is reflected by the high relative abundances of many effector/immunity and T6SS^iii^ markers compared to other diagnoses (Figure 1A). Much lower median profile alpha diversity was observed in infant microbiomes than in adults, but with high interindividual variability. This finding raises hypotheses that polymorphic toxins may play a role in microbiome development, building diversity from infancy to adulthood. Effector/immunity profile beta diversity also readily distinguished microbiomes by disease state (Bray-Curtis dissimilarity adonis2 p < 0.001) (Figure 1C).

### Polymorphic toxin effector/immunity profiles predict disease states

The diversity patterns within the complete meta-analysis dataset suggested that effector/immunity profiles may be predictive of disease state. However, differences in study design and methodology may also contribute to variance. For example, depth of sequencing may influence alpha diversity, and effector/immunity profiles may be affected by factors such as geographic location and population demographics. To address this possibility, we compared beta diversity by diagnosis within each study (as defined in Table S2). The primary metrics applied were pairwise differences in Bray-Curtis dissimilarity measurements within the diagnostic group and across diagnostic groups and permutation-based statistical testing with adonis2 (Figure 1D). All diagnoses had distinguishable effector/immunity profiles within individual studies, except for antibiotics-treated *vs.* healthy controls. The profiles of each diagnosis also had significant beta diversity differences from the meta-analysis pooled healthy controls and compared to the entire dataset (Figure 1D). Finally, we validated the correlation between PolyProf diversity and disease state using an independent cohort of village residents (Figure S5)^1^. The findings indicate that human diseases have distinct effector/immunity patterns that can be identified within a large population of subjects of diverse health statuses.

We next leveraged the large dataset to generate machine learning predictors of IBD based on polymorphic effector/immunity profiles. We built predictive models for IBD (by combining CD or UC diagnoses) and comparing to “healthy” specimens or to all non-IBD specimens in the studies defined in Table S2 (Figure 1E). Using patient-grouped, 10-fold cross validation, we optimized the XGBoost hyperparameters and tested the model on a 20% hold-out dataset. Polymorphic effector/immunity profiles were predictive of IBD, with predictive model performance meeting or exceeding current models built using taxonomy ^40^. Predictive model performance (hold-out balanced accuracy 0.86) increased with sample size in the IBD vs. all other diagnoses dataset, overcoming the heterogeneity introduced by other disease-related dysbiosis (Figure 1E).

We next applied the IBD model hyperparameters to train new predictive models for each of the other dataset diagnoses (Figure 1E). Polymorphic effector/immunity profiles were highly predictive across the range of diseases. In general, the diagnosis *vs.* all other metagenomes models outperformed the diagnosis *vs.* healthy control models, reflecting improved performance with larger training datasets. Particularly high-performing models (testing area under the receiver operating curve [AUC] and balanced accuracy >= 0.85) were identified for osteoarthritis, rheumatoid arthritis, type 2 diabetes, smoking, IBD, obesity, antibiotic use, and metabolic syndrome. Taken together, the polymorphic toxin effector/immunity profiles show strong potential for predicting disease-specific dysbiosis.

Since PolyProf and taxonomy are not highly interdependent, we reasoned that combined microbiome relative abundance and PolyProf data would likely enhance model performance. We generated XGBoost models with hyperparameter tuning to distinguish each individual diagnosis from all others. Combined taxonomy and PolyProf model performance on hold-out validation datasets was near perfect, with AUC 1.00 and balanced accuracy 0.97-1.0 for all 12 diagnoses depicted in Figure 1E.

### PolyProf correlates with markers of active inflammation in IBD

Since PolyProf is predictive of disease state, we hypothesized that disease-related markers may correlate with disease severity. A subset of 662 IBD metagenomes had accompanying fecal calprotectin measurements, a marker of neutrophilic inflammation in active disease ^41^. Multiple PolyProf marker abundances showed linear correlation with fecal calprotectin concentrations (Figure S6). Among these were inverse correlations with Tde, a marker whose depletion we previously associated with IBD ^25^, and the predicted RNase Toxbarnase, the top distinguishing feature in the machine learning predictor for IBD diagnosis (Figure S10). We conclude that PolyProf is likely related to disease severity in IBD. Active inflammation is accompanied by depletion of markers that are predictive of an IBD diagnosis.

### Infant delivery mode shapes the microbiome toxin secretion system and effector/immunity profiles

Caesarean section delivery (CSD) markedly affects the intestinal microbiome composition of infants during the first year of life ^13–15,20^. However, at 1 year and beyond, microbiomes of CSD and vaginally delivered (SVD) babies tend to converge and resemble the maternal microbiome in terms of species diversity ^12,14,42^. We hypothesized that polymorphic toxin secretion systems may be particularly active in the developing microbiome, modulating colonization by new strains through interbacterial antagonism. We tested this hypothesis by looking for PolyProf profile changes over time, stratified by birth delivery mode.

We first examined PolyProf global diversity as a function of infant age and delivery mode (Figure 2). Like previous descriptions of taxonomy ^12,14,42^, polymorphic toxin effector/immunity profiles of infants evolve through the first year of life with a tendency toward the maternal PolyProf. PolyProf beta diversity is strongly influenced by the delivery mode, with a closer similarity to maternal PolyProf in vaginally delivered infants (Figure 2A). Analysis of paired maternal-infant metagenomes ^14,43^ showed a similar trend of gradual PolyProf convergence toward the maternal microbiome over the first year of life, quantified as decreasing Bray-Curtis dissimilarity of infant and maternal microbiomes as a function of age (Figure 2B, Skillings-Mack test p < 0.001). Acquisition of new and increasing diversity of polymorphic toxin effector/immunity markers is reflected as an early life increase in PolyProf alpha diversity (Figure 2C, Skillings-Mack test p < 0.001). Using an independent dataset from families in Guinea-Bissau ^44^, we compared PolyProf beta diversity distances by family relations (Figure S7). Parent-child PolyProf were more similar within families than in unrelated adult-child pairs. Child-child and adult-adult distances did not differ by family status. The same pattern was observed in a large USA-based cohort of oral (saliva) microbiomes (Figure S7) ^44^. Specifically, mother-child and not father-child or mother-father PolyProf distances were shorter within families than unrelated pairs. Oral and fecal microbiome PolyProf differ substantially, reflected by highly separate clustering on beta diversity NMDS (not shown). The findings are consistent with vertical transmission, primarily of maternal origin, resulting in fecal and oral PolyProf similarities between related parents and children.

**Figure 2.**
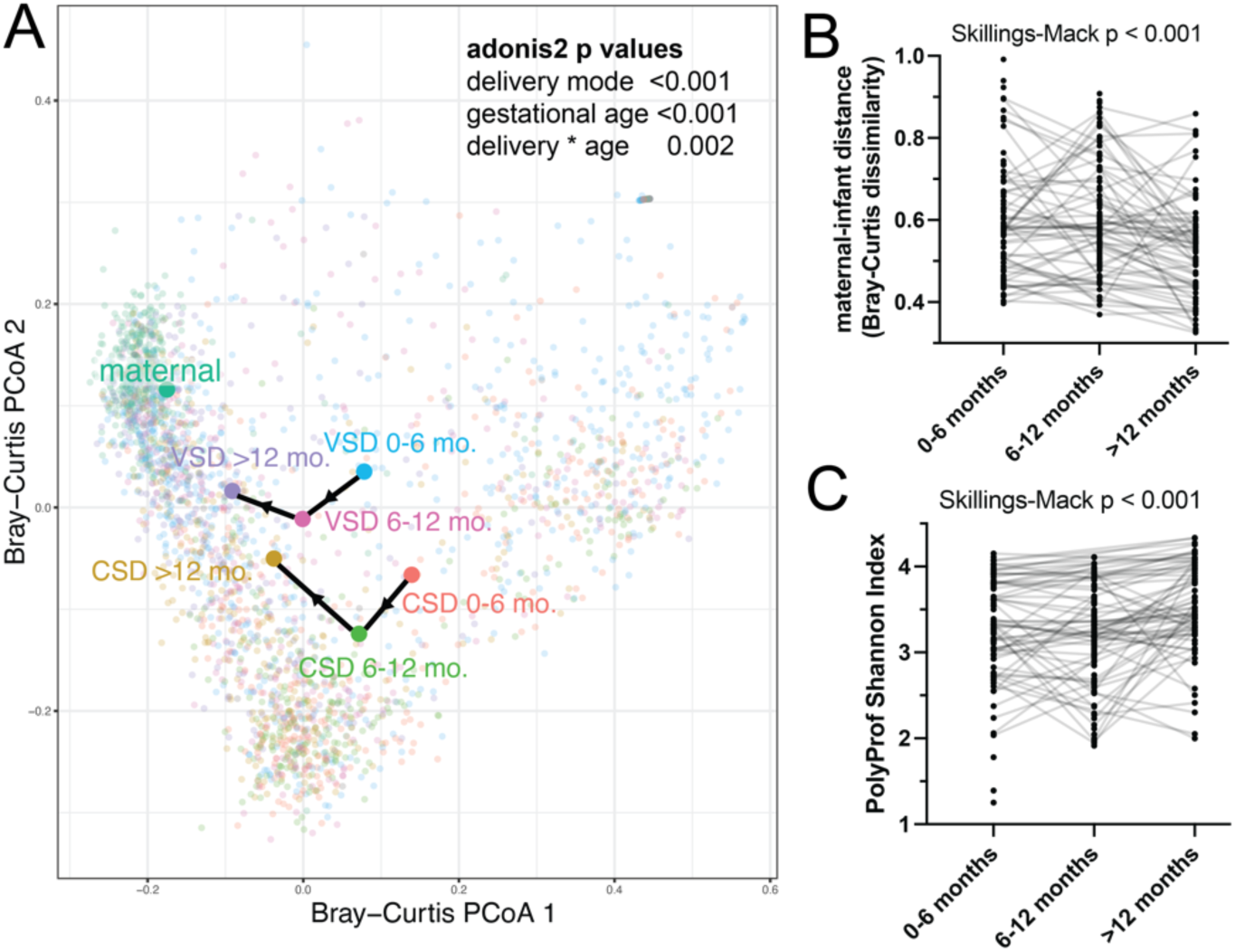
Infant microbiome polymorphic toxin effector/immunity profiles develop toward maternal profiles in the first year of life. (A) PolyProf beta diversity of infant microbiomes is distinct from maternal microbiomes. Infant PolyProfs are significantly affected by delivery mode (VSD vaginal delivery or CSD Cesarean section). As infants age, PolyProf becomes more similar to maternal microbiomes, shown by centroids (large circles) and time course with black lines. For subjects with known maternal-infant pairings (connected with lines), the trend toward infant similarity with the maternal PolyProf over time is reflected as decreasing Bray-Curtis dissimilarity between maternal and infant PolyProfs (B). PolyProf alpha diversity also increases with age (C).

We next asked which specific markers underly the delivery mode-dependent PolyProf diversity shifts in the first year of life. In infant biospecimens collected between 6 and 12 months after delivery, there was strong enrichment of T6SS^iii^ and several effector/immunity markers in SVD, and this pattern persisted beyond 1 year of life (Figure 3A,B). PolyProf markers showed disparate dependence on delivery mode and gestational age (Figure 3C). T6SS^iii^ is highly related to vaginal delivery, while the predicted RNase Ntox26 abundance is most strongly related to the subject’s age. The findings indicate interplay of several factors in determining the toxin secretion system and effector/immunity profiles of developing microbiomes.

**Figure 3.**
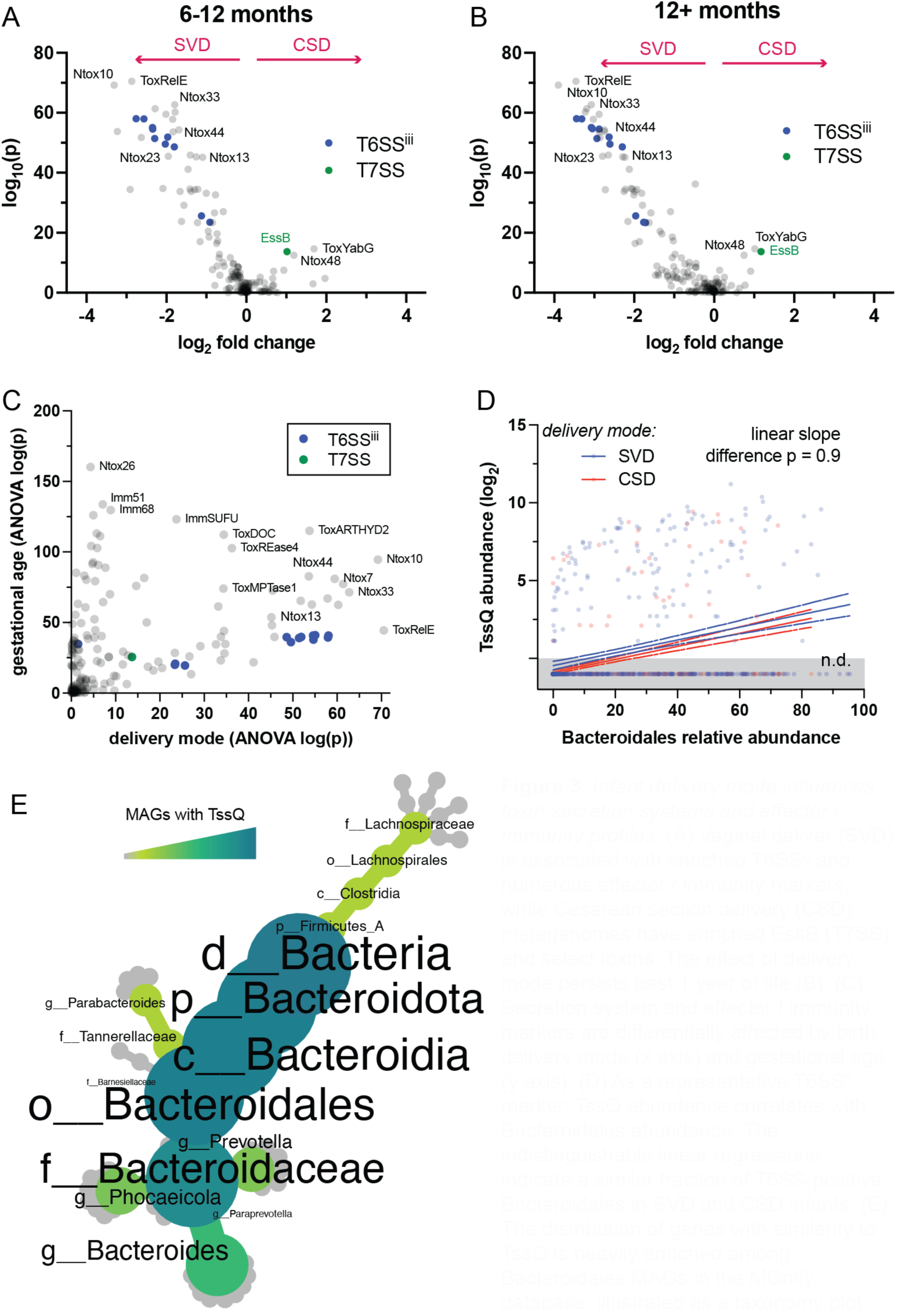
Infant delivery mode influences toxin secretion systems and effector/immunity profiles. (A) Vaginal deliver (SVD) is associated with enriched T6SS^iii^ and numerous effector/immunity markers, while Cesarean section delivery (CSD) metagenomes have enriched EssB (T7SS) and select toxins. The effect of delivery mode persists past 1 year of life (B). (C) Secretion system and effector/immunity markers are differentially affected by birth delivery mode (x axis) and gestational age (y axis). (D) As a representative T6SS^iii^ marker, TssQ abundance correlates with Bacteroidales abundance. The indistinguishable linear regressions indicate a similar fraction of T6SS-positive Bacteroidales in SVD and CSD infants. (E) The distribution of genes with similarity to TssQ is heavily enriched among Bacteroidales MAGs in the MGnify database, illustrated as a taxonomy plot.

Since T6SS^iii^ is highly restricted to order Bacteroidales, we asked whether the delivery mode dependence of this secretion system is simply reflecting differential abundances of Bacteroidales (Figure 3D,E). Depleted Bacteroidales were a prominent feature of CSD microbiomes in prior studies ^13,14^. As expected, the abundance of T6SS^iii^ markers (TssQ shown as a representative) is related to Bacteroidales abundance determined using MetaPhlAn4 ^38^. The relationship is not linear because a minority of Bacteroidales strains carry T6SS^iii^. In fact, many metagenomes had Bacteroidales without any detectable TssQ (Figure 3D). Linear regression was used to estimate the fraction of T6SS^iii^-positive Bacteroidales, which was not distinguishable by delivery mode. The findings indicate that CSD infants have depleted gut microbiome T6SS^iii^ systems by virtue of lower abundances of Bacteroidales.

### Maternal effector/immunity repertoires influence newborn microbiome development

Prior research indicates that as newborn microbiomes develop through the first year of life, they tend to become increasingly like the maternal microbiome in terms of taxonomic diversity ^14^. There is also strong evidence for maternal-infant transmission of bacterial strains both at delivery and throughout the first year of life ^15,16,18^. Figure 2 shows a similar pattern of PolyProf convergence with the maternal microbiome over time. We hypothesized that transmission of polymorphic effector/immunity and secretion system encoding strains from mother to infant influences microbiome development, and we predicted that the presence of these markers in the maternal microbiome would strongly predict detection in the infant. To address this, we focused on infants with paired maternal metagenomes ^14,43^.

T6SS^iii^ systems and several effector/immunity markers in infant metagenomes were strongly related to their detection in the maternal metagenomes (Figure 4A). TssQ as a representative of T6SS^iii^ was more abundant in metagenomes of infants born to mothers whose microbiomes also carry TssQ, and this difference increased through the first year of life (2-way ANOVA p < 0.001, Figure 4B). A higher fraction of Bacteroidales carried TssQ in infants born to mothers with detectable TssQ than those without, illustrated as a linear slope difference (p < 0.001 Figure 4C). Bacteroidales were more abundant in the metagenomes of infants with likely vertical transmission (TssQ positive mother and infant) than those without (Figure 4E). This finding suggests that encoding T6SS^iii^ confers an advantage to Bacteroidales over other taxonomic groups. Other effector/immunity markers such as Imm12 show a similar pattern of higher abundance in infants whose mother’s microbiome carries the same gene (Figure 4D). Imm12 immunity is frequently encoded adjacent to and predicted to protect against Tox-URI2 effectors, predicted DNases ^3^. Imm12 and other effector/immunity markers correlating to maternal detection are not restricted to Bacteroidales. The findings provide strong evidence for maternal–infant transmission of polymorphic secretion system and effector/immunity encoding bacteria. The time course patterns suggest that transmission continues through the first year of life.

**Figure 4.**
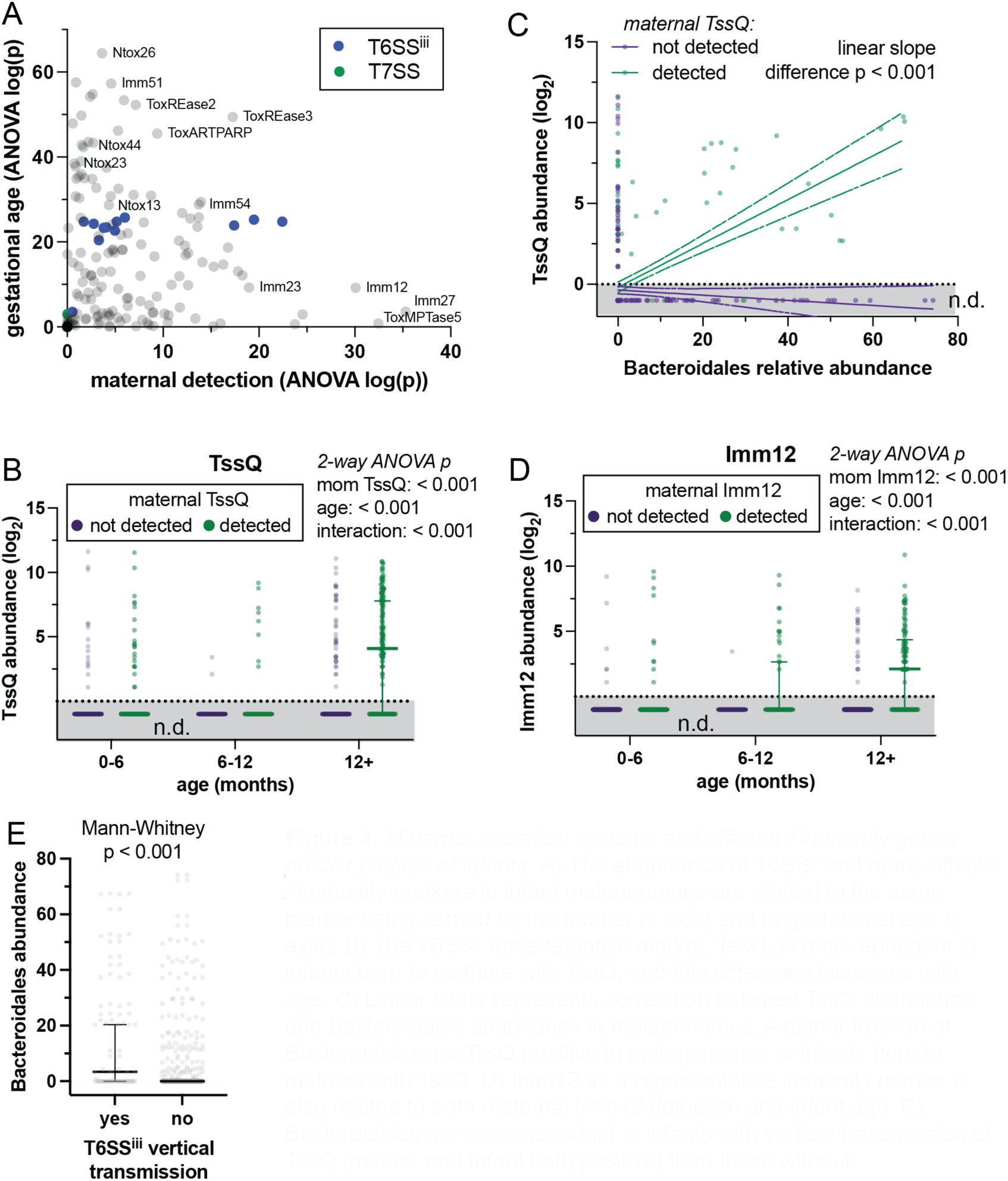
Maternal secretion systems and effector/immunity genes predict profiles of infants. A) The abundance of T6SS^iii^ and many effector/immunity markers in infant metagenomes are related to the same marker being carried by the mother (x axis) and to gestational age (y axis). B) The T6SS^iii^ representative marker, TssQ, is more abundant in infants born to mothers with TssQ, and this difference increases with age. C) Linear slope represents correlation between TssQ abundance and Bacteroidales abundance in metagenomes. A higher fraction of Bacteroidales are TssQ positive in metagenomes of infants born to mothers with TssQ. D) Imm12 as a representative immunity marker is also related to both maternal Imm12 detection and infant age. E) Bacteroidales are more abundant in infants with vertical transmission of TssQ (mother and infant both positive) than those without.

### Breastfeeding changes effector/immunity profiles

Secretion system and effector/immunity encoding strain transmission from mother to infant continues to occur after SVD, and prior research has shown that breastfeeding is an important mode of microbiome vertical transmission ^18^. We asked whether breastfeeding status also affects PolyProf profiles in the infant microbiome. To address this, we focused analysis on SVD infant metagenomes stratified by breastfeeding status ^14,43,45^.

The T7SS structural protein EssB and several effector/immunity markers were significantly associated with breastfeeding status and gestational age (Figure 5A). Ntox26 is shown as an example effector that was lower abundance in exclusively breastfed infants than in non-exclusively breastfed or unknown status infant microbiomes (Figure 5B). Both Ntox26 and EssB (T7SS) are predominantly distributed among Bacillota MAGs. We conclude that breastfeeding also impacts infant PolyProf profiles during the first year of life, although the effects are smaller in magnitude than delivery mode and maternal profiles.

**Figure 5.**
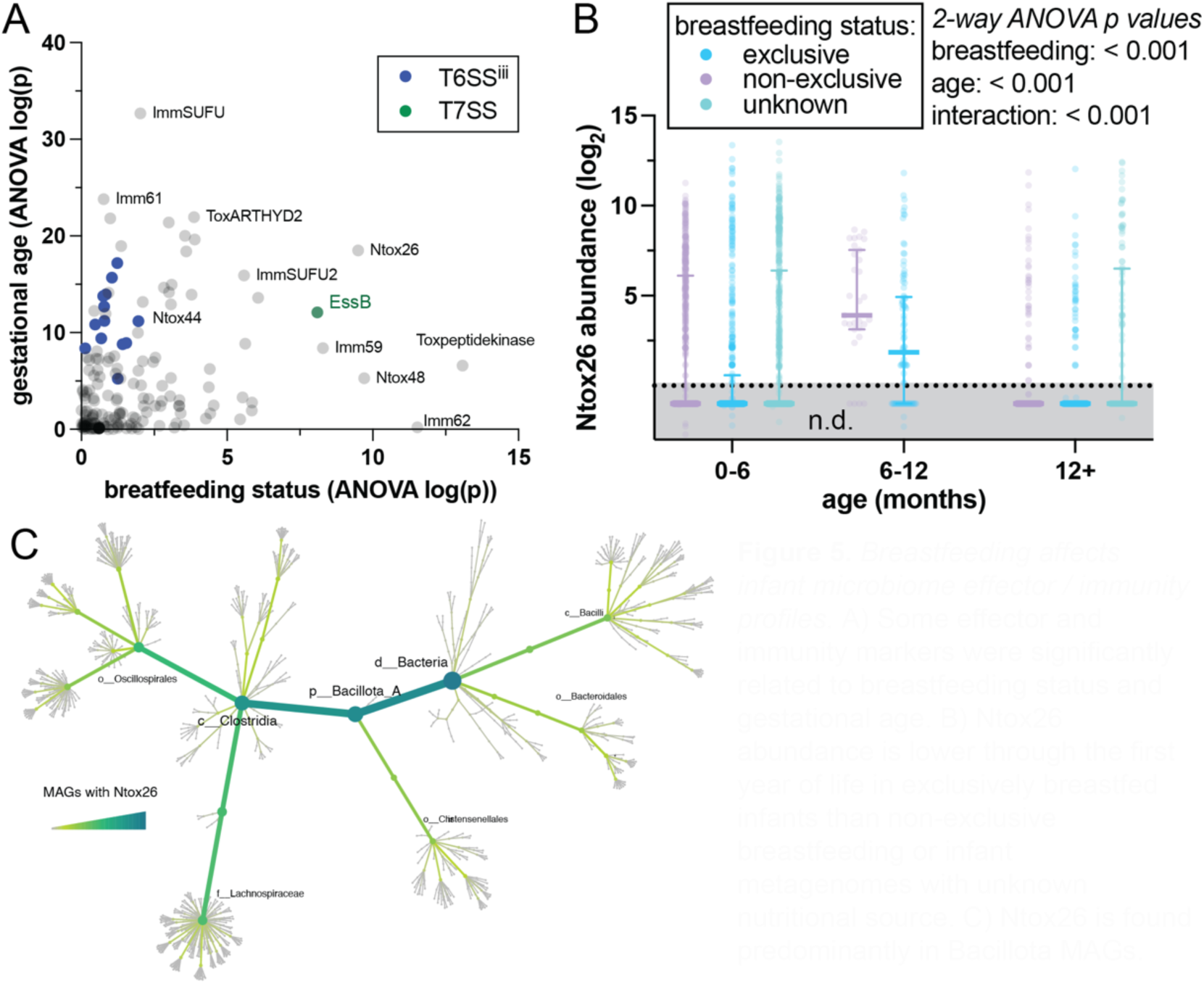
Breastfeeding affects infant microbiome effector/immunity profiles. A) Some effector and immunity markers were significantly related to breastfeeding status and gestational age. B) Ntox26 abundance is lower through the first year of life in exclusively breastfed infants than non-exclusive breastfeeding or infant metagenomes with unknown nutritional sources. C) Ntox26 is found predominantly in Bacillota MAGs.

### Effector/immunity profiles are related to social strain sharing

Close contact is likely an essential requirement for vertical strain transmission. Recent evidence suggests that gut bacterial strains are also shared during social interactions ^1^. We hypothesized that polymorphic toxin effector/immunity encoding strains may have a competitive advantage in establishing colonization during social contact strain sharing. If the hypothesis is correct, we expected that PolyProf would be more similar among socially connected individuals and correlate with strain sharing.

To test these hypotheses, we applied PolyProf to 1117 fecal metagenomes from subjects living in isolated villages in Honduras ^1^. PolyProf alpha and beta diversity clustered by village (de-identified indicators), and high strain sharing rates between individuals correlated with more similar PolyProf (Figure 6). Co-residency in the same village or in the same building had strong and independent correlation with beta and alpha diversity (Figure 6A,B). PolyProf beta diversity was most similar (shortest Bray-Curtis dissimilarity distance) for subject pairs within a family, followed by non-family social contacts, and village co-residents (Figure 6C). Subject pairs in all social relation groups had more similar PolyProf when strain sharing was detected (Figure 6C, 2-way ANOVA p < 0.001). In the whole dataset, shorter PolyProf beta diversity pairwise distances correlated with higher strain sharing rates. We conclude that PolyProf similarity is highly correlated with strain sharing among social contacts. Although the strongest effects are seen for family and close social contacts, PolyProf patterns are distinguishable on the village level. Thus, social context is a key determinant of PolyProf on the population (village-wide) level.

**Figure 6.**
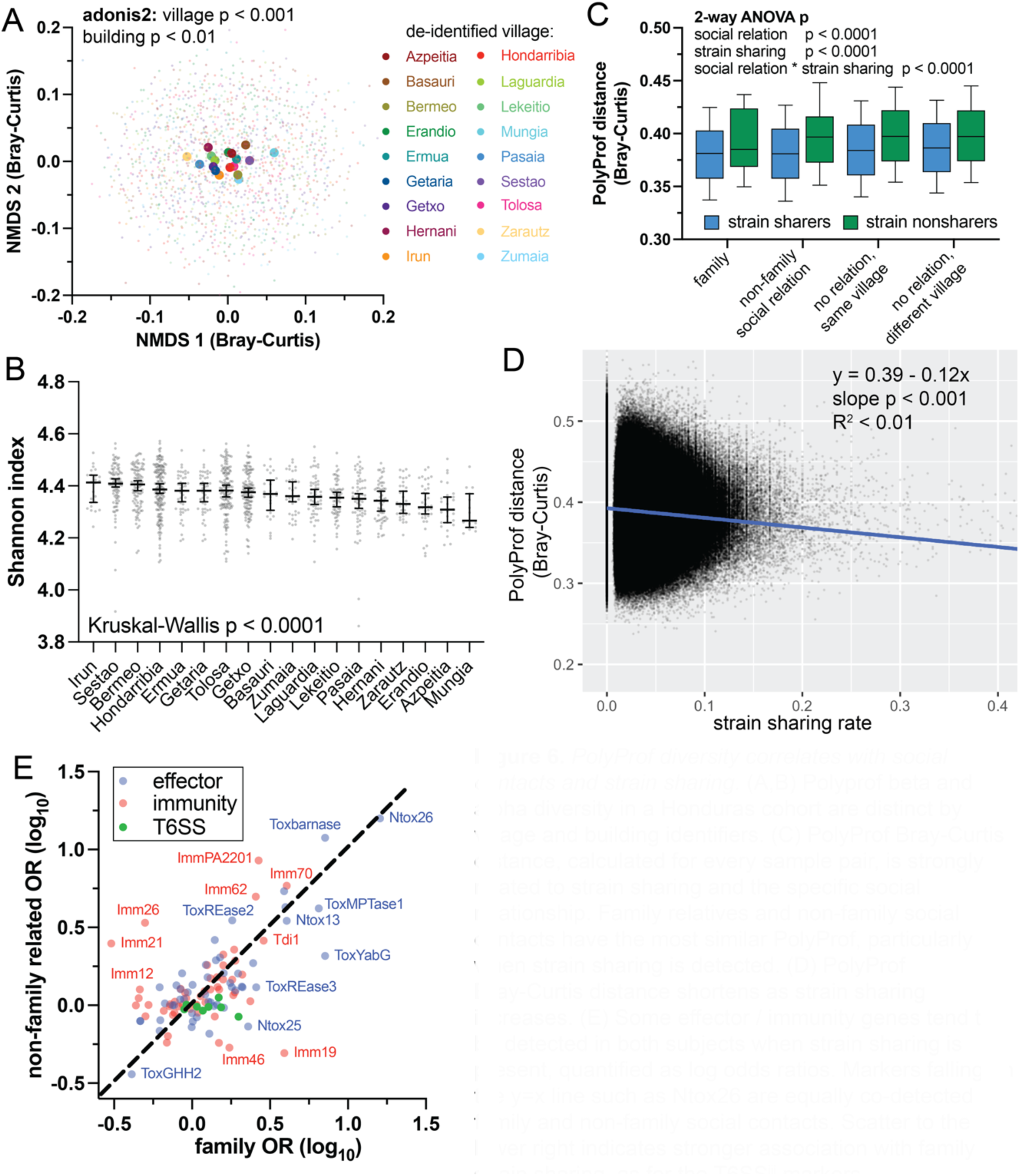
PolyProf diversity correlates with social contacts and strain sharing. (A,B) Polyprof beta and alpha diversity in a Honduras cohort are distinct by village and building identifiers. (C) PolyProf Bray-Curtis distance, calculated for every sample pair, is strongly related to strain sharing and the specific social relationship. Family relatives and non-family social contacts have the most similar PolyProf, particularly when strain sharing is detected. (D) PolyProf Bray-Curtis distance shortens as strain sharing increases. (E) Some effector/immunity genes tend to be detected in both subjects when strain sharing is present, quantified as log odds ratios. Markers falling on the y=x line such as Ntox26 are equally co-detected in family and non-family social contacts. Scatter to the lower right indicates a stronger association with family strain sharing, as for the T6SS^iii^ markers.

We next asked which specific effector/immunity genes are most highly involved in social strain sharing. We reasoned that marker genes promoting strain transmission will be more likely co-detected in subject pairs that share strains (donor and recipient). From 2×2 tables of effector/immunity co-detection and strain sharing detection, odds ratios were calculated and tested with Fisher’s exact statistics (Figure 6E). Most effector/immunity markers were either neutral (OR ∼1) or more frequently co-detected in strain sharers (OR > 1). Some markers were co-detected in strain sharers with specific types of social interaction. For example, Imm 21 co-detection is enriched in non-family social contacts but not families. Consistent with the prior mother-infant data, T6SS^iii^ promotes strain sharing within families (reflecting vertical transmission), but does not drive non-family strain sharing (co-detection ORs ∼1, Figure 6E). Since all subjects in the cohort are adults, the results indicate persistence of T6SS^iii^ effects on transmission beyond infancy and childhood. The most highly strain sharing-associated marker Ntox26 was also prominently associated with vertical transmission during microbiome development (Figures 3-5), collectively implicating this effector as a likely important driver of strain transmission in multiple contexts.

To identify strain-level PolyProf and secretion system features related to strain sharing, we assembled 130,852 MAGs from Honduras cohort metagenomes and calculated their relative transmissibility between individuals. By applying TXSScan ^46^ to all MAGs and associating detection of secretion system genes with transmissibility, we identified two systems positively associated with transmission: Tad and flagellum (Figure S11-S12). These two systems have diverse known functions that include but are not limited to effector secretion. A Tad-like system in *Myxococcus xanthus* mediates contact-dependent interbacterial antagonism ^47,48^. The flagellar type III secretion system exports proteins for flagellum assembly and is also known to secrete antimicrobial toxins in some strains ^49^. Our approach is limited in determining which functions of these systems (*e.g.* adhesion, motility, or secretion) are involved in strain transmission. Three specific effector classes, Ntox13, Ntox23, and Ntox44 (all three predicted RNases) were enriched in highly transmissible MAGs, detected with gene set enrichment analysis (Figure S13). All three effectors were identified in MAGs of diverse taxonomy and had poor overlap with Tad and flagellum systems. All 3 effector types were also significantly associated with infant microbiome development and vertical transmission (Figures 3-5), implicating these effectors as a likely important drivers of strain transmission in multiple contexts.

## Discussion

Here, we have developed and applied a marker sequence-based approach to profile polymorphic toxin secretion systems and effector/immunity classes in gut metagenomes. These systems are known to mediate interbacterial antagonism within the intestinal ecosystem ^28,34^, but little was previously known about their relationship to dysbiosis and human disease. A central finding of this study is that PolyProf readily distinguishes adult human disease states, which indicates that polymorphic toxins are an integral component of dysbiosis. We speculate that many of the observed associations are likely effects of the disease state rather than causes. For example, intestinal inflammation in IBD drastically changes the bacterial microenvironment with the release of oxygen species, mucosal disruption and bleeding, and compromise of mucin barriers ^10^. In the dysbiotic state, interbacterial interactions are also perturbed, which may result in competitive advantages (or disadvantages) of certain effectors and corresponding immunity proteins. Our study highlights several disease-associated markers that suggest avenues for mechanistic studies in disease models to address the directionality of causation. The performance of machine learning models based solely on PolyProf is comparable to similar approaches using bacterial taxonomy ^40^. Combining PolyProf and bacterial taxonomy enabled the development of near-perfect predictive models. This finding emphasizes the importance of strain-level functional characteristics like secretion systems in disease dysbiosis that are not fully accounted for with traditional metagenomic taxonomy analyses. There is a substantial promise for enhanced disease-related classifiers that incorporate the quantitation of known functionally important genes in combination with taxonomy.

Our second major finding is that polymorphic toxin secretion systems and effector/immunity profiles are highly dynamic and variable in the developing microbiome of infants. Exposure to maternal bacteria at delivery (vaginal vs. surgical), vertical strain transfer, and breastfeeding status all shape the PolyProf profile diversity through the first year of life. In one example, we show that the presence of T6SS^iii^ can influence the abundance of Bacteroidales and play a role in shaping microbiome taxonomic diversity. As Verster *et al.* have modeled and demonstrated experimentally in *Bacteroides fragilis*, T6SS^iii^ effectors mediate antagonism in the intestine and exclude related, non-immune Bacteroidales from colonization ^37,50^. Thus, vertical transmission of bacteria with secretion systems and immunity to the dominant effectors is likely one mechanism by which developing newborn microbiomes gravitate toward the parental microbiome, excluding competing bacteria to which the infant is exposed.

Our study focused on the dominant role of maternal-infant transmission because of data availability, but recent studies indicate stable transmission from paternal sources as well ^17^. We raise the hypothesis that vertical polymorphic effector immunity transmission plays a role in the nongenetic inheritance of disease. Seminal studies in the microbiome of obesity, for example, have demonstrated that the transfer of the microbiome from obese individuals to mice is sufficient to promote weight gain ^7^. Does the dysbiotic microbiome and PolyProf of a parent with obesity get transmitted to offspring and increase the risk for obesity later in life?

Roles for polymorphic toxin secretion systems in microbiome development are not restricted to the first year of life. Our third major finding is that PolyProf is related to strain sharing during social interactions. Beghini et al. showed that gut microbiome strain sharing among a cohort of people living in isolated villages in Honduras occurs through both familial and non-familial social interactions ^1^. We find that PolyProf beta diversity and specific effector/immunity genes are in turn related to strain sharing, not only in families, but also with close social contacts and village co-residents. These combined studies indicate ongoing exchange of commensal bacteria through human social interactions after the first year of life. Some polymorphic toxin secretion system effector and immunity genes may even influence strain sharing, likely through competitive colonization mechanisms.

## Conclusions

PolyProf accurately quantifies polymorphic toxin secretion effector/immunity genes in shotgun metagenomics datasets, distinguishes many disease states, and can be used to construct high-performing decision tree machine learning models to predict disease. Polymorphic toxin secretion effector/immunity genes are involved in parent-child vertical strain transmissions and strain sharing among non-family social contacts. In summary, this study adds correlational evidence that polymorphic toxin secretion systems, effectors, and immunity proteins are important determinants of initial microbiome development and social strain sharing, and they are markers of diverse adult disease statuses. Further study is needed to determine their mechanistic roles in disease and for the development of potential microbiome-based diagnostics.

### Limitations

The results and conclusions of the meta-analysis are somewhat dependent on the accuracy of the metadata. “Healthy control” designations are particularly study specific. For example, the “healthy controls” of an IBD study may include tobacco users and confound the smokers vs. all healthy controls analysis. However, the expected effect would be increased variance in the control group and bias toward the null hypothesis (smokers and healthy controls not distinguishable) and therefore lead to *underestimation* of disease-specific effects and lowered performance of disease classifiers. As in all meta-analyses, population and methodologic differences could affect the cross-study comparisons. While marker sequences are a well-established method for gene family quantitation, off-target mapping could increase the variance and bias toward the null hypothesis for any marker family. The power and accuracy of machine learning models are dependent on sample size, uniformity, and specific parameters employed. The parameters used in this study were optimized for IBD and then applied to all other comparisons for uniformity and comparability. Better-performing models are certainly achievable, even simply by testing alternative algorithms and diagnosis-specific hyperparameter tuning.

## Methods

### Data and software resources

Previously published and publicly available shotgun metagenomic data from human fecal specimens were used for PolyProf validation and meta-analysis (Table S2) ^11,14,19,43,45,51–73^. Taxonomic profiling was performed with Kraken 2 ^74^. Polyprof profiling was performed with a custom protein sequence marker database and HUMAnN 3 ^38^. The effector/immunity sequence database was constructed using NCBI BLAST ^75^ and HMMER ^76^, with the MGnify gut metagenome-assembled genome (MAG) database ^77^. Queries were representative sequences and hidden Markov models from each polymorphic toxin effector and immunity class described in the Zhang et al. comprehensive bioinformatic analysis ^3^. Also included were Bacteroidales type VI secretion system (T6SS^iii^) conserved structural proteins and the conserved type VII secretion system (T7SS) structural protein EssB. A stringent sequence similarity cutoff was selected to minimize off-target mapping of metagenomic reads (i.e. high specificity). The tradeoff is the potential lack of detection of divergent members of the effector/immunity classes (low sensitivity). Enrichment of each effector/immunity marker for each taxonomic lineage was calculated based on the fraction of positive MAGs within the lineage and corresponding sampling error. Enrichment analyses for all markers are available on the PolyProf GitHub page. Taxonomic trees for each marker were visualized using the R *metacoder* package ^78^.

The marker sequences were compiled into a custom HUMAnN database using DIAMOND ^79^. The marker sequence database is not intended to be exhaustive, as other effector types are known to be secreted through various secretion systems and mediate interbacterial antagonism. However, the PolyProf database used in this study can readily be expanded to include other effector/immunity marker sequences of interest. Other published resources and databases are excellent resources for identifying other effector/immunity classes ^80–83^. The PolyProf effector/immunity database and source sequences will be made publicly available on the PolyProf GitHub page. Abundance tables for all metagenome profiles as part of this study will also be available.

Marker gene performance testing was performed by simulating Illumina-style microbiome data using InSilicoSeq ^39^. One million reads were simulated from random selections of 10 known marker-positive MAGs and 100 marker-negative MAGs. Simulation was repeated 5 times for each marker. PolyProf was applied to simulation fastq files, and marker abundance correlated to marker-positive MAG abundance using linear regression (GraphPad Prism). 1:1 correlation of these abundances was expected and observed. Deviation of PolyProf marker quantities from the value expected based on MAG abundance was measured for each simulation with mean and 95% confidence interval calculation over the five replicates to identify markers that were systematically over- or under-estimated by PolyProf.

Contributions of metagenome taxonomy to PolyProf variance were estimated with two approaches using the IBD study set. Taxonomic abundances were calculated with MetaPhlAn4 ^84^. NMDS analysis (R vegan and phyloseq packages) was performed separately on both the taxa relative abundances and PolyProf, and the dominant NMDS factor values were plotted for each sample. Linear regression was performed and R^2^ calculated as an estimate of how much PolyProf variance is explained by taxonomic beta diversity. The second approach to assess each individual marker’s relationship to taxonomy was to predict marker abundance from family-level taxonomy and the rate of marker detection in MAGs from each family: *predicted abundance = family detection rate in MAGs * family relative abundance in the metagenome*, summed for all families. This taxonomy-predicted abundance was compared to the PolyProf marker abundance using linear regression and R^2^ calculation. Graphs for each marker with linear regression data are available on the PolyProf GitHub site. Contributions of metagenome Bacteroidales abundances to PolyProf associations with diagnoses were assessed using MaAsLin3 multivariable linear modeling of all PolyProf markers by diagnosis, with and without adjustment for Bacteroidales order abundances ^85^.

### Disease-related dysbiosis comparisons

PolyProf profiling was performed on 14,740 human gut metagenomes using HUMAnN 3 ^38^ and default settings other than input of the PolyProf custom protein database and bypass of the nucleotide search. Non-detection was assigned a value of 0.5 for downstream statistical testing. Polymorphic effector/immunity beta diversity was compared within each study set as defined in Table S2. Exceptions were osteoarthritis, obesity, and rheumatoid arthritis studies with did not include non-disease control groups. Individual disease-distinguishing markers were identified using multivariate logistic regression of marker relative abundances with diagnosis as the dependent variable. Beta diversity was measured as Bray-Curtis dissimilarity and alpha diversity as Shannon index using the *vegan* R package. The meta-analysis design before beginning analysis was to perform 3 tiered comparisons: 1) compare diagnosis groups within each study, 2) compare diagnosis groups to all “healthy controls”, and 3) compare diagnosis groups to all other metagenomes. Each comparison was Shannon index alpha diversity with Mann-Whitney or Kruskal Wallis tests as appropriate, Bray-Curtis beta diversity distance testing with adonis2 (*vegan* R package), and LefSe linear discriminant analysis ^86^. Global beta diversity was visualized using NMDS analysis of Bray-Curtis distance matrices. All pairwise comparison statistical tests were adjusted for FDR 0.01 using the Benjamini-Hochberg method. Within the IBD cohort, 662 patients had available fecal calprotectin data. Calprotectin was linearly correlated with each PolyProf marker using the *corr* package in R.

### Machine learning classifier generation and testing

Binary supervised machine learning models were trained to predict diagnoses with two tiers of polymorphic effector/immunity comparisons: diagnosis *vs*. healthy subjects across all metagenome samples utilized in this study (Table S2), and diagnosis *vs*. all other metagenomes. Each sample consisted of 217 effector or immunity gene target hits from the custom PolyProf HUMAnN database (in reads per kilobase, adjusted for length). For combined taxonomy and PolyProf models, the dataset also included ∼5,000 bacterial relative abundances. To prevent data leakage among individual subjects, individuals within longitudinal studies were assigned a host ID and grouped together for model cross-validation, training, and testing. Models were trained using 80% of individuals and tested on a holdout dataset of 20% of individuals. The testing and training datasets were transformed by removing the mean and scaling to a variance of one for each variable, with parameters for scaling determined from the training dataset. Initial cross-validated training indicated that the XGBoost algorithm had the highest balanced accuracy among a variety of algorithms tested (not shown), and XGBoost was selected for further predictive model development. Hyperparameter tuning was conducted using “IBD” as the diagnosis, which was a combination of “UC” or “CD” diagnoses. Hyperparameter tuning was then conducted with a 10-fold subject-grouped cross-validation on the training set (full details on hyperparameters tested and selected are shown in Table S3). As an initial demonstration of PolyProf predictive potential and to minimize training time, optimal hyperparameters identified from training on “IBD” diagnoses were utilized to train models for the alternative diagnoses. For each alternative diagnosis from “IBD,” the only parameter changed between models was the positive weight scaling, which was set to [sum(negative instances) / sum(positive instances)] in the training set, to account for unbalanced class distribution. The XGBoost algorithm^87^ was implemented in Python (v. 3.11.4) using the *xgboost* module (v. 2.1.1) and data preparation and model metrics were calculated with the Scikit-Learn module (v. 1.1.3)^88^. Predictive models were evaluated by calculating and plotting the area under the receiver operating curve (AUC), balanced accuracy, and plotting the AUC and the balanced accuracy.

### Infant delivery mode, maternal-infant transmission, and breastfeeding status microbiome analysis

PolyProf profiling and taxonomic quantitation with MetaPhlAn4 ^84^ were applied to all infant and maternal samples. For delivery mode analyses, only infant microbiomes with metadata indicating delivery mode were included. Differences in marker gene abundances were calculated as log_2_-fold change using median values and pairwise statistical testing using Student’s t-tests, visualized as a volcano plot. Effects of delivery mode and gestational age were measured using ANOVA. Infant ages were binned as 0-6 months, 6-12 months, or >12 months. Marker sequence changes were quantified over these age bins and stratified by delivery mode. Statistical testing was a 2-way ANOVA. For maternal-infant transmission studies, only infant metagenomes with a linked maternal metagenome ^14^ were included. Any non-zero marker gene quantity in the maternal metagenome was defined as detection. Time course paired analysis of PolyProf diversity was tested using a generalization of the Friedman test for missing data called the Skillings-Mack test ^89^.

### Social strain sharing microbiome analysis

One study ^1^ included potentially identifying metadata with protected access. For this study, human subjects research was conducted in accordance with the Declaration of Helsinki, with Iowa IRB protocol (#202406297). The original dataset was collected with informed consent of the participants, and this study is a secondary use of the data. PolyProf was applied to metagenome data from a Honduras cohort which had strain sharing scores previously calculated ^1^. PolyProf beta diversity was compared by resident village and building identifiers using adonis2 and 2-way ANOVA testing of Bray-Curtis dissimilarities. Pairwise beta diversity distances were correlated with strain sharing rates using linear regression. For each marker, a 2×2 table of strain sharing (zero or not zero) and marker co-detection (present in both subject’s metagenomes or not) was constructed. Odds ratios were calculated and p-values derived by Fisher’s exact tests.

### Metagenome assembled genomes

Metagenomic assembly was performed on all samples using metaSPADES ^90^ (v. 3.13.1, default parameters), and contigs longer than 1000bp were discarded. Reads were mapped against the assembled contigs using minimap2 ^91^ (v. 2.24, parameters: -ax sr -t 12 -N50). The resulting alignments were used as input for the VAMB pipeline for contig binning ^92^ (v. 3.0.9, parameters: --minfasta 500000). The automatic binning procedure generated a total of 226,039 MAGs that were then subjected to quality control to evaluate completeness and contamination using CheckM2 ^93^ (v. 1.0.2). We selected a total of 130,852 MAGs passing the thresholds for medium-quality genomes, being at least 50% complete and displaying less than 10% of contamination. All MAGs were annotated with Bakta ^94^ (v. 4.1, default parameters) using the “Full” database. MAGs were dereplicated at 95% ANI using galah ^95^, which yielded a total of 2,609 metagenomic species clusters. Cluster representatives were taxonomic annotated with the phylophlan_assign_sgb routine from PhyloPhlAn 3 (v. 3.1.1) ^96^ using the SGB Jan 21 database. ^97^

### MAG transmissibility, TXSS, and PolyProf analysis

Previously calculated strain sharing scores were used to calculate species-level transmissibility, which is the number of strain-sharing events detected for a species divided by the total number of potential strain-sharing events for all the species detected by StrainPhlAn4. MAG representatives for each species were used for downstream analysis and transmissibility values were then linked to them. Predicted proteins were used as input for MacSyFinder2 ^98^ to detect components of secretion systems with the TXSScan ^46^ using the unordered search mode. In addition to the detection of secretion systems with TXSScan, we aligned the previously built database of effector/immunity sequences against all the predicted proteins using JACKHMMER. The resulting alignment was filtered to retain only domain hits with e-value <0.001 and domain coverage of at least 50%. Species were ranked according to their transmissibility value and for each effector, enrichment analysis was performed using gene set enrichment analysis as implemented in the fgsea R package ^99^, by using species with the effector detected as present as gene set. Phylogenetic trees were generated using the Newick tree file provided by MetaPhlAn 4 ^84^ and annotated with ggtree ^100^.

## Data and code availability statement

Source metagenomic data is publicly available from the Sequence Read Archive and accession numbers are given in Table S2. The PolyProf database, sequences, and associated graphical data are available on GitHub at https://github.com/dustin-bosch/PolyProf/.

## Author contributions

HWS – conceptualization, methodology, investigation, resources, writing (original draft and editing), visualization; FB – conceptualization, methodology, investigation, visualization; JARG – methodology, software, formal analysis, investigation, resources, data curation, writing (review and editing); NAC – resources, supervision; DEB – conceptualization, methodology, investigation, software, resources, data curation, writing (original draft and editing), visualization, supervisions, project administration, funding acquisition.

## Acknowledgements

DEB was supported by the NIH, K08AI159619.

---

**Table S1.** *Marker genes included in PolyProf.*

**Table S2.** *Shotgun metagenomics studies included in the meta-analysis*. NCBI BioProject accessions, disease state and case number summaries, and corresponding prior publications are listed.

**Table S3.** *XGBoost hyperparameters tested and selected using grid search and leave-one-subject-out cross validation.*

**Figure S1.** *Polymorphic toxin effector/immunity profiling meta-analysis design.* Fecal metagenomic sequence data was profiled using a custom HUMAnN database of MAG-derived effector/immunity protein sequences. Effector/immunity abundance profiles were analyzed to identify disease-specific associations.

**Figure S2.** *Co-abundance correlations of all marker genes across metagenome datasets.* Marker genes are clustered according to similarity. Color and ellipse shape indicate the degree of linear least squares abundance correlation. T6SS^iii^ markers exhibit high linearity of co-abundance, reflecting genetic co-occurrence in T6SS loci which require all 13 structural components for assembly of the secretion apparatus.

**Figure S3.** *PolyProf marker performance on simulated metagenomic data.* Metagenomic data were simulated from MAGs with and without known corresponding effector/immunity genes. Strong linear correlation between simulated taxon abundance and PolyProf marker abundance was observed (black linear regression line with dashed 95% confidence intervals). Six markers had marker abundance measurements that deviated significantly from the expected value based on taxon abundance, highlighted with colored diamonds.

**Figure S4.** *PolyProf variance is weakly related to taxonomic beta diversity.* Metagenomes (n = 1537) from the IBD cohort including healthy controls were analyzed with PolyProf and MetaPhlAn, followed by Bray-Curtis dissimilary NMDS analysis. (A) The dominant PolyProf and taxonomy NMDS values for each metagenome are fitted with linear regression. The R^2^ value of 0.11 indicates that approximately 11% of PolyProf variance is explained by taxonomic beta diversity. (B) Linear regression models for each marker comparing taxonomy-based predicted and actual PolyProf-measured abundances (plots available on GitHub). Variance explained by taxonomy (R2) is shown for the 10 most highly taxonomy-correlated markers. Ntox4 abundance is exceptionally predictable from taxonomy, followed by eight components of the Bacteroidales-restricted T6SS^iii^. (C) Associations of all PolyProf markers with disease state were assessed with and without adjustment for Bacteroidales order abundance using MaAsLin3 ^85^. The model coefficient represents strength and direction (negative = inverse relationship) of abundance associations. Some T6SS^iii^ associations are impacted by adjustment for Bacteroidales, but remain statistically significant. (D) A similar plot of only T6SS^iii^ markers, stratified by disease state, demonstrates the highest taxonomy dependence of T6SS^iii^ associations with infant microbiomes and rheumatoid arthritis. Other diseases like UC have a T6SS^iii^ association that is essentially independent of Bacteroidales abundance.

**Figure S5.** *PolyProf diversity correlates with disease state in an independent Honduras cohort.* A) Alpha diversity was compared by participant self-reported disease, symptoms, or tobacco use. Obesity status was assigned based on an objectively measured BMI > 30 kg/m2. Diarrhea, allergies, and smoking were associated with significant alpha diversity deviation from healthy controls (* p < 0.05 or ** p <0.01, adjusted for multple comparisons). B) PolyProf beta diversity also differed by disease, with strongest contributions from smoking, obesity, and diabetes in multivariate adonis2 testing.

**Figure S6.** *Correlation of fecal calprotectin with PolyProf.* (A) Fecal calprotectin was linearly correlated with all PolyProf markers in 662 participants with available data. (B, C) Top correlating markers ToxREase2 and Imm62 had decreased abundance in patient with active inflammation (increased fecal calprotectin). p values represent slope differences from zero. Calprotectin also inversely correlated with Tde (D), a previously published marker depleted in IBD, and Toxbarnase (E), the top distinguishing feature in IBD machine learning classifiers (Figure S10).

**Figure S7.** *Fecal and oral microbiome PolyProf are similar in related mother-child pairs.* (A) In a fecal microbiome dataset from a Guinea-Bissau population, PolyProf beta diversity differences measured by Bray-Curtis dissimilarity were lower in related parent-child pairs. Between-child and -adult distances were not different according to family relationships. (B) Oral microbiomes from a USA cohort were most similar in related mother-child pairs. Boxplots represent median and interquartile range. *** p < 0.001 Mann-Whitney test. n.s. not significant.

**Figure S8.** *ROC curves for decision tree models of each disease vs. all healthy controls.*

**Figure S9.** *ROC curves for decision tree models of each disease vs. all other diagnoses.*

**Figure S10.** *Top distinguishing features for decision tree models of each disease vs. all other diagnoses.* Importance of features in XGBoost models was quantified with permutation-based testing. The top 10 features for each diagnosis are shown.

**Figure S11.** *Honduras cohort Tad secretion system association with MAG transmissibility.* All metagenome-assembled genomes were compared with a phylogenetic tree (center). Presence or absence of Tad system genes as detected with TXSScan are shown in yellow and purple, respectively. MAG transmissibility (outer ring) is depicted by quartile. Tad is enriched and associated with transmission in Lachnospiraceae and Bacilli.

**Figure S12.** *Honduras cohort flagellum system association with MAG transmissibility.* All metagenome-assembled genomes were compared with a phylogenetic tree (center). Presence or absence of flagellum system genes as detected with TXSScan are shown in yellow and purple, respectively. MAG transmissibility (outer ring) is depicted by quartile. Flagella are enriched and associated with transmission in Lachnospiraceae.

**Figure S13.** *Polymorphic effector associations with MAG transmissibility.* All metagenome-assembled genomes were compared with a phylogenetic tree (center). Presence or absence of three transmissibility-enriched effectors as detected with hmmer are shown in yellow and purple, respectively. MAG transmissibility (outer ring) is depicted by quartile. All 3 transmission-associated effectors are dispersed across diverse taxa.

