## Supplemental Figures for "Metagenomic polymorphic toxin effector and immunity profiling predicts microbiome development and disease-related dysbiosis"

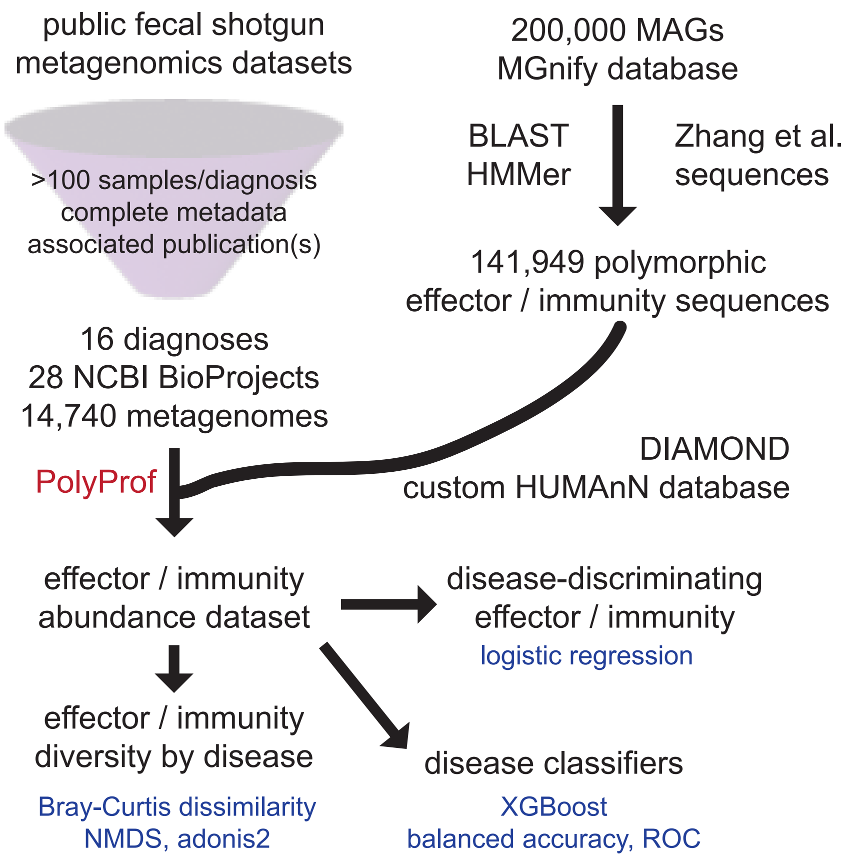


**Figure S1.** *Polymorphic toxin effector / immunity profiling meta-analysis design.* Fecal metagenomic sequence data was profiled using a custom HUMAnN database of MAG-derived effector / immunity protein sequences. Effector / immunity abundance profiles were analyzed to identify disease-specific associations.

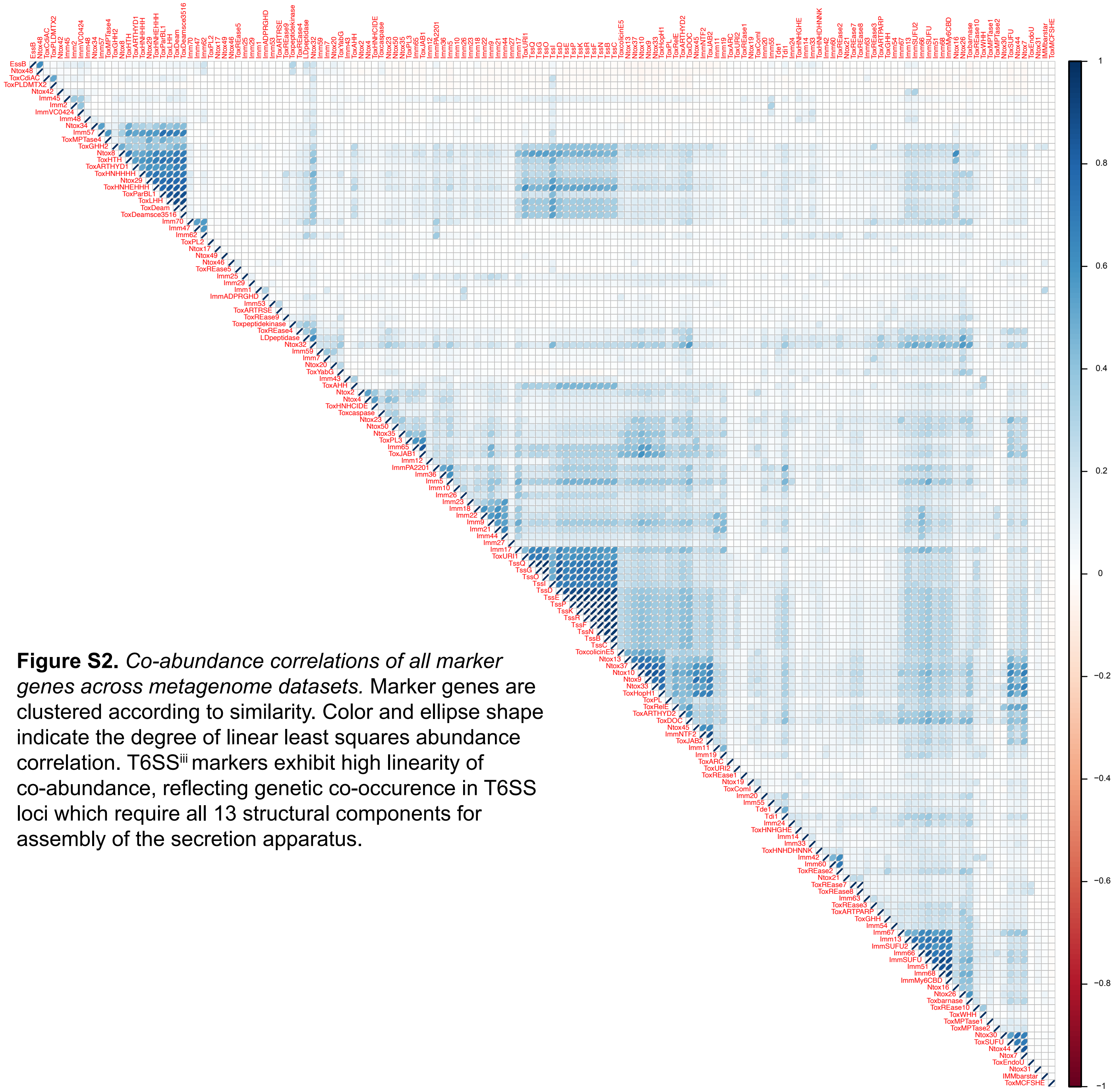


**Figure S2.** *Co-abundance correlations of all marker genes across metagenome datasets.* Marker genes are clustered according to similarity. Color and ellipse shape indicate the degree of linear least squares abundance correlation. T6SS$^{iii}$ markers exhibit high linearity of co-abundance, reflecting genetic co-occurence in T6SS loci which require all 13 structural components for assembly of the secretion apparatus.

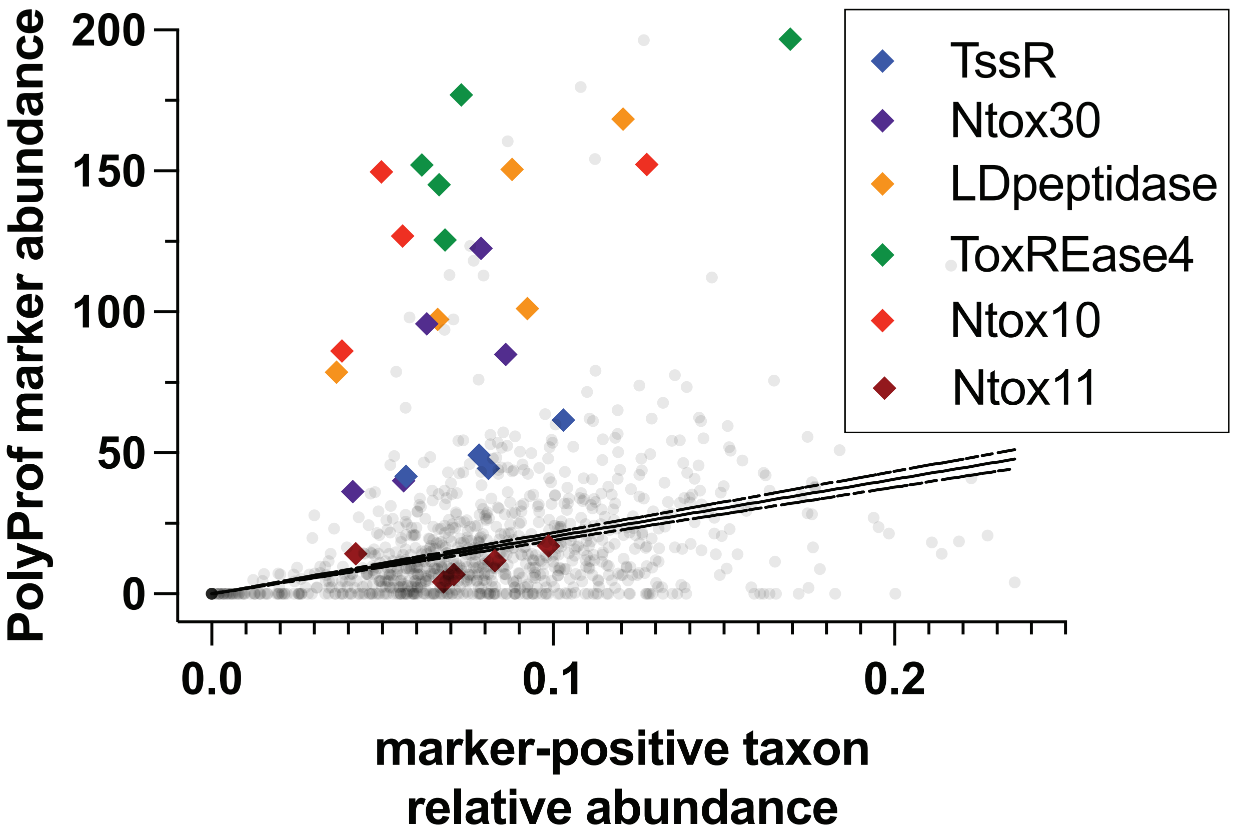


**Figure S3.** *PolyProf marker performance on simulated metagenomic data.* Metagenomic data were simulated from MAGs with and without known corresponding effector / immunity genes. Strong linear correlation between simulated taxon abundance and PolyProf marker abundance was observed (black linear regression line with dashed 95% confidence intervals). Six markers had marker abundance measurements that deviated significantly from the expected value based on taxon abundance, highlighted with colored diamonds.

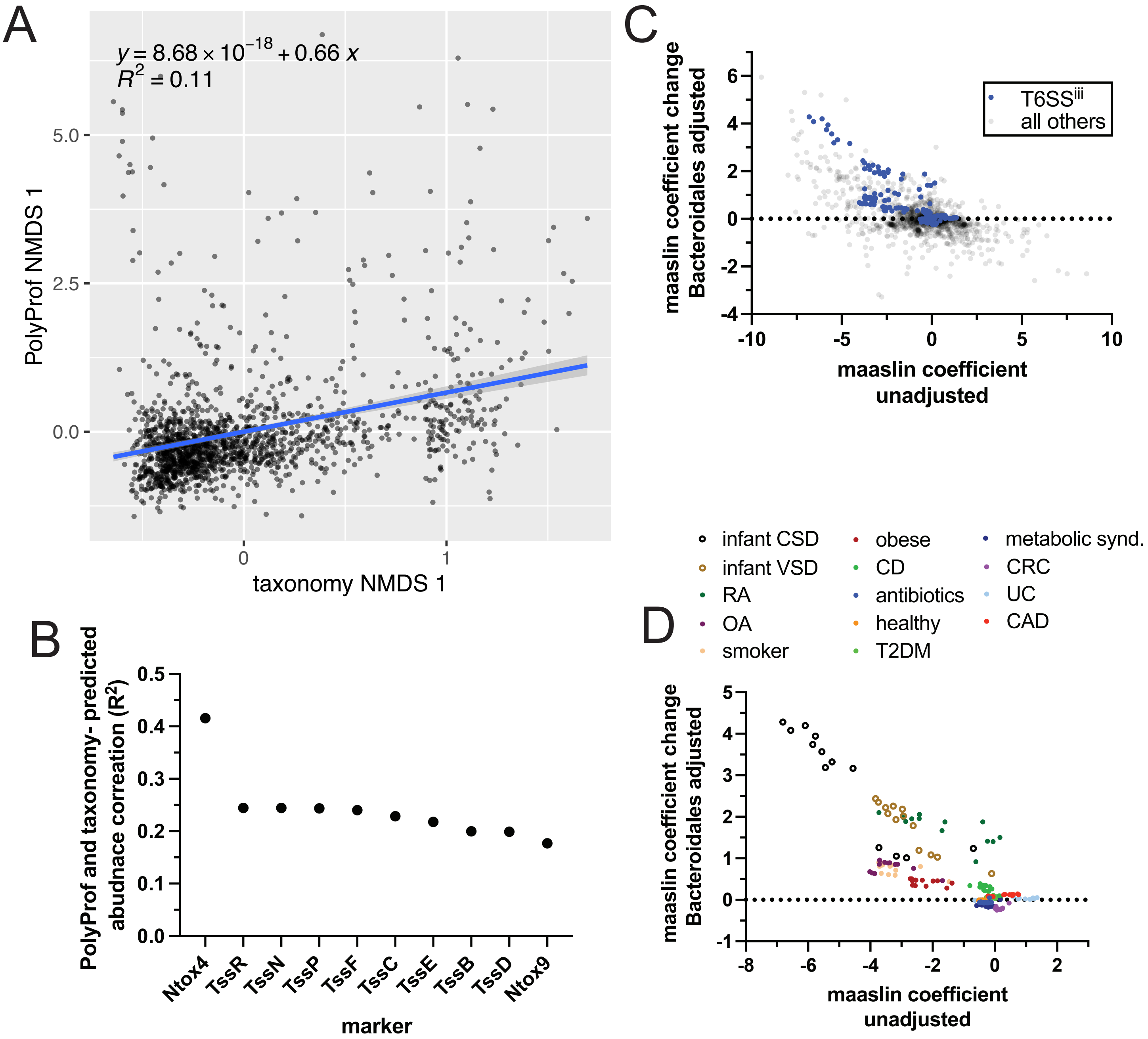


**Figure S4.** *PolyProf variance is weakly related to taxonomic beta diversity.* Metagenomes (n = 1537) from the IBD cohort including healthy controls were analyzed with PolyProf and MetaPhlAn, followed by Bray-Curtis dissimilary NMDS analysis. (A) The dominant PolyProf and taxonomy NMDS values for each metagenome are fitted with linear regression. The $R^2$ value of 0.11 indicates that approximately 11% of PolyProf variance is explained by taxonomic beta diversity. (B) Linear regression models for each marker comparing taxonomy-based predicted and actual PolyProf-measured abundances (plots available on GitHub). Variance explained by taxonomy ($R^2$) is shown for the 10 most highly taxonomy-correlated markers. Ntox4 abundance is exceptionally predictable from taxonomy, followed by eight components of the Bacteroidales-restricted $T6SS^{iii}$. (C) Associations of PolyProf marker abundances with diagnoses were calculated using MaAsLin3, with and without adjustment for Bacteroidales abundance. The differences in model coefficients upon Bacteroidales adjustment are plotted vs. the unajusted coefficients. A y-axis value of zero would indicate no effect of Bacteroidales abundance on PolyProf-disease associations. Some $T6SS^{iii}$ diagnosis associations are influenced by taxonomy. (D) To identify specific diagnoses where $T6SS^{iii}$ abundance is driven by taxonomy, a similar plot shows only $T6SS^{iii}$ maker associations, colored by diagnosis. For some populations, such as infants (open circles) $T6SS^{iii}$ is highly affected by Bacteroidales abundance. Other diagnoses, such as UC and CAD, have$T6SS^{iii}$ maker enrichment that is largely independent of Bacteroidales abundance.

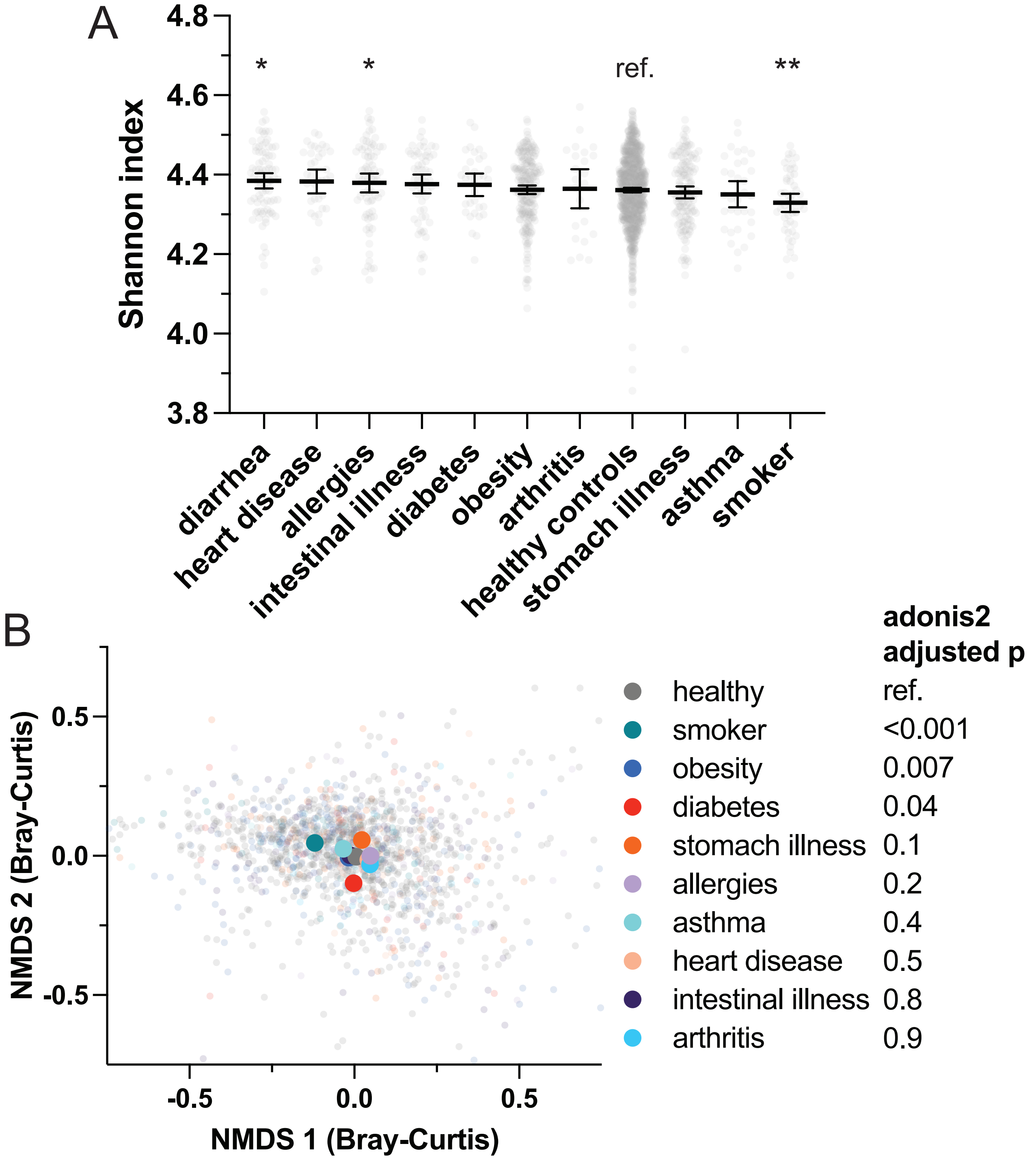


**Figure S5.** *PolyProf diversity correlates with disease state in an independent Honduras cohort.* A) Alpha diversity was compared by participant self-reported disease, symptoms, or tobacco use. Obesity status was assigned based on an objectively measured BMI > 30 kg/m$^2$. Diarrhea, allergies, and smoking were associated with significant alpha diversity deviation from healthy controls (* p < 0.05 or ** p <0.01, adjusted for multple comparisons). B) PolyProf beta diversity also differed by disease, with strongest contributions from smoking, obesity, and diabetes in multivariate adonis2 testing.

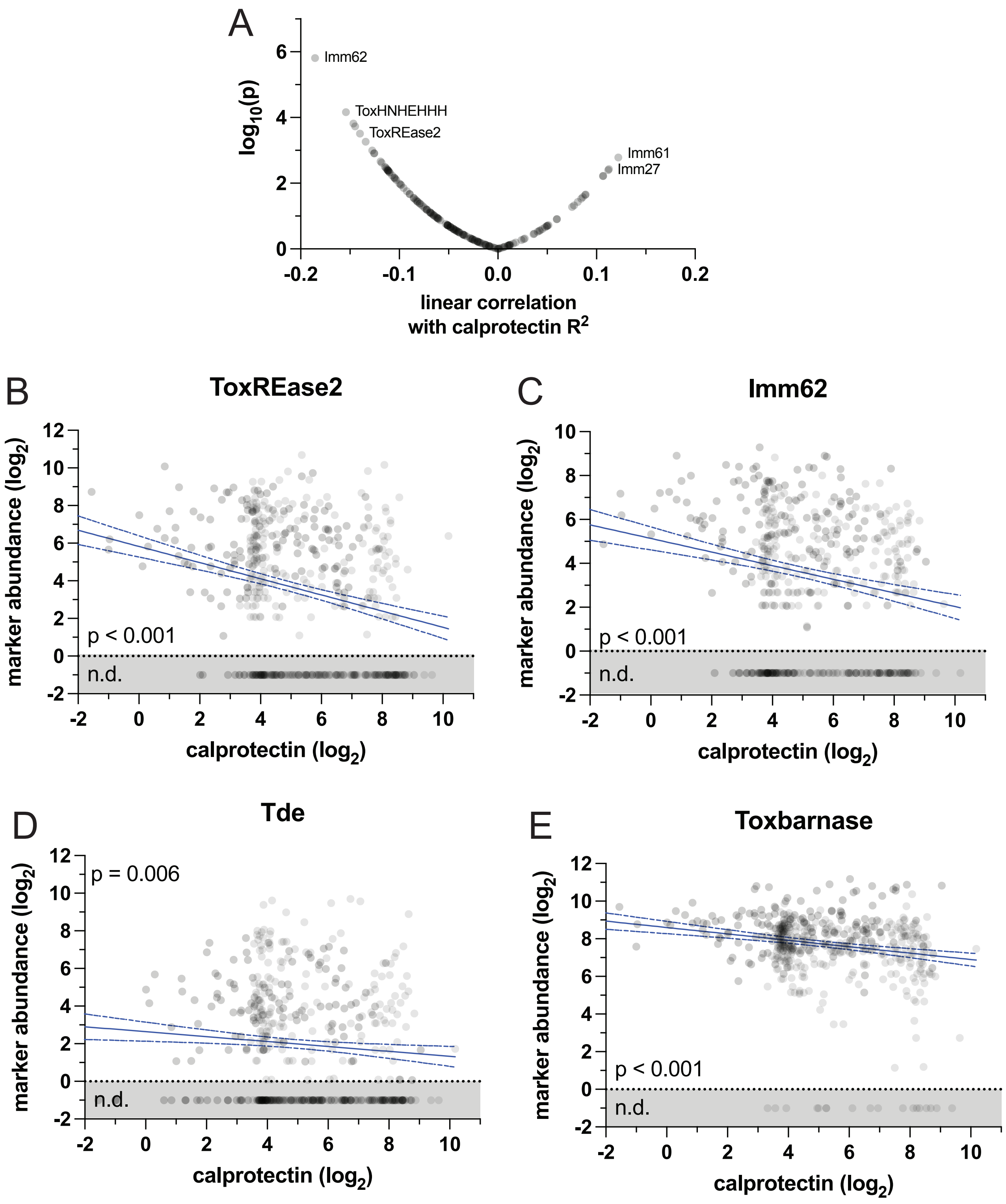


**Figure S6.** *Correlation of fecal calprotectin with PolyProf.* (A) Fecal calprotectin was linearly correlated with all PolyProf markers in 662 participants with available data. (B, C) top correlating markers ToxREase2 and Imm62 had decreased abundance in patient with active inflammation (increased fecal calprotectin). p values represent slope differences from zero. Calprotectin also inversely correlated with Tde (D), a previously published marker depleted in IBD, and Toxbarnase (E), the top distinguishing feature in IBD machine learning classifiers (Figure S9).

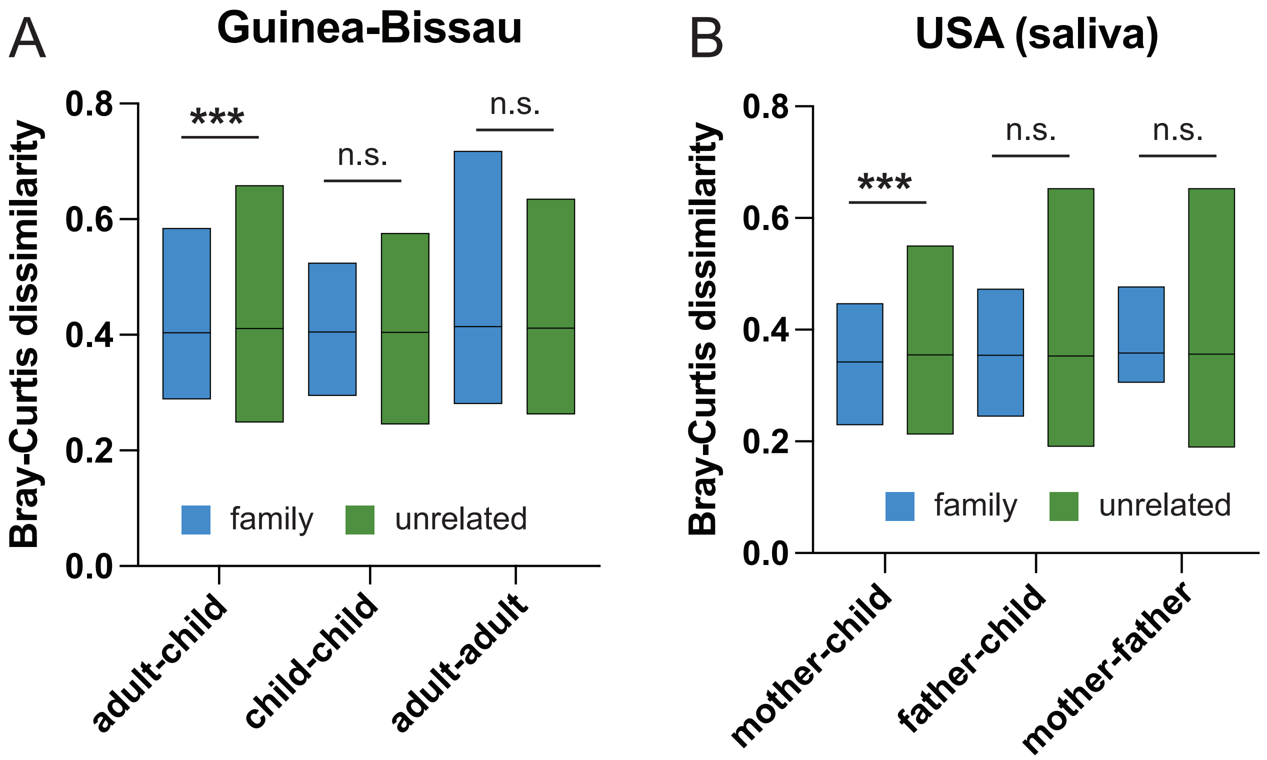


**Figure S7.** *Fecal and oral microbiome PolyProf are similar in related mother-child pairs.* (A) In a fecal microbiome dataset from a Guinea-Bissau population, PolyProf beta diversity differences measured by Bray-Curtis dissimilarity were lower in related parent-child pairs. Between-child and -adult distances were not different according to family relationships. (B) Oral microbiomes from a USA cohort were most similar in related mother-child pairs. Boxplots represent median and interquartile range. *** $p < 0.001$ Mann-Whitney test. n.s. not significant.

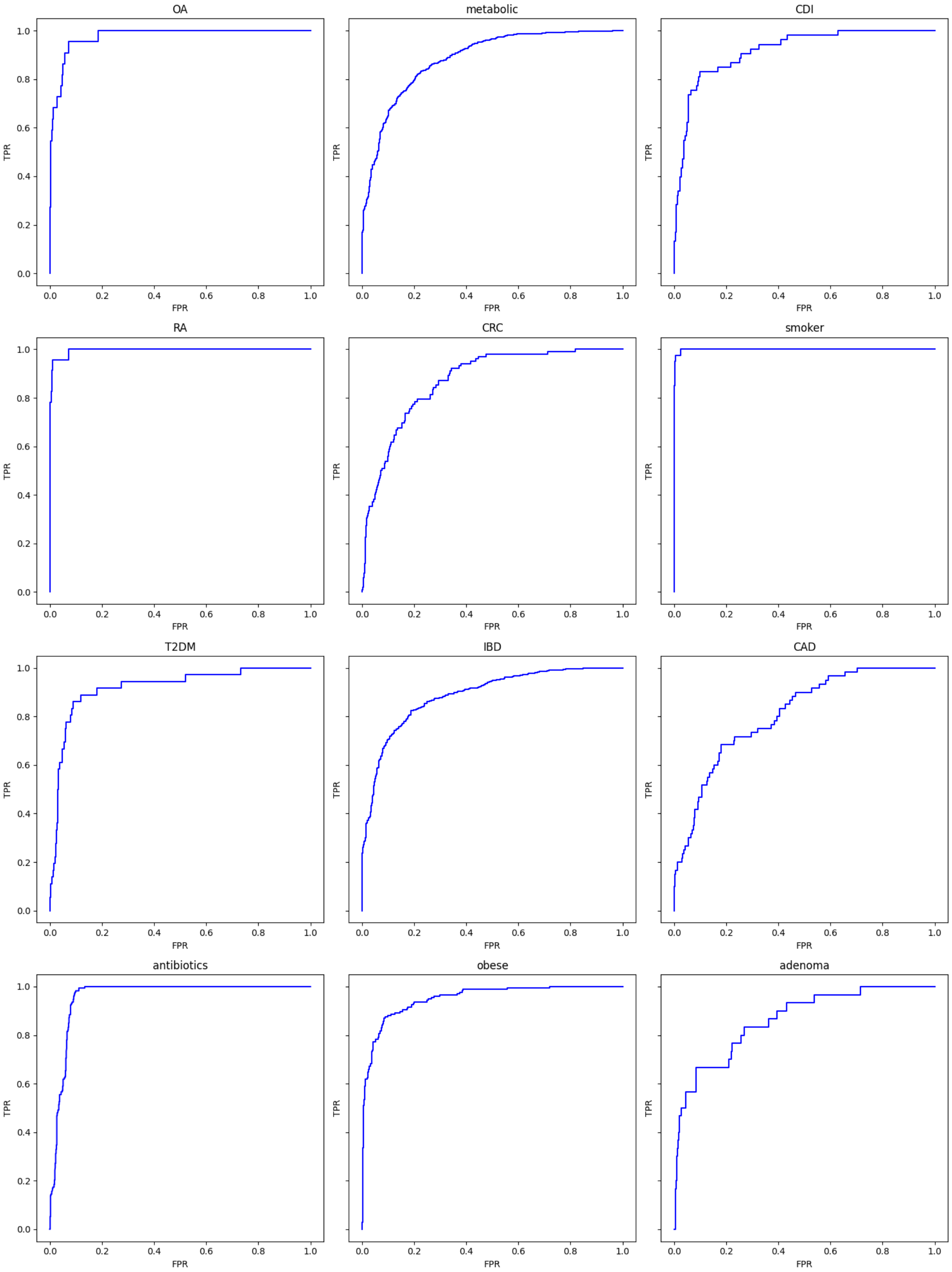


**Figure S8.** *ROC curves for decision tree models of each disease vs. all healthy controls.*

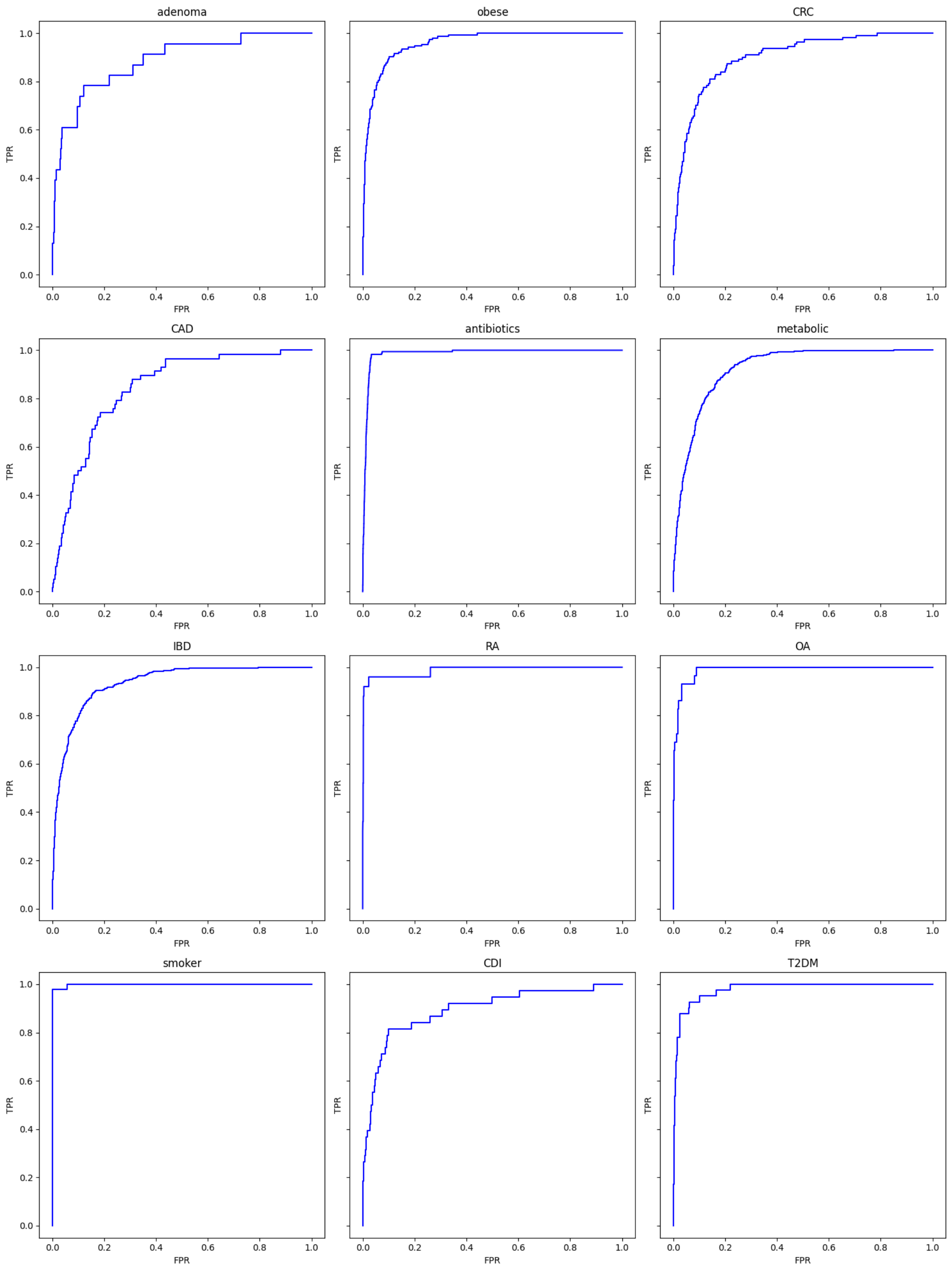


**Figure S9.** *ROC curves for decision tree models of each disease vs. all other diagnoses.*

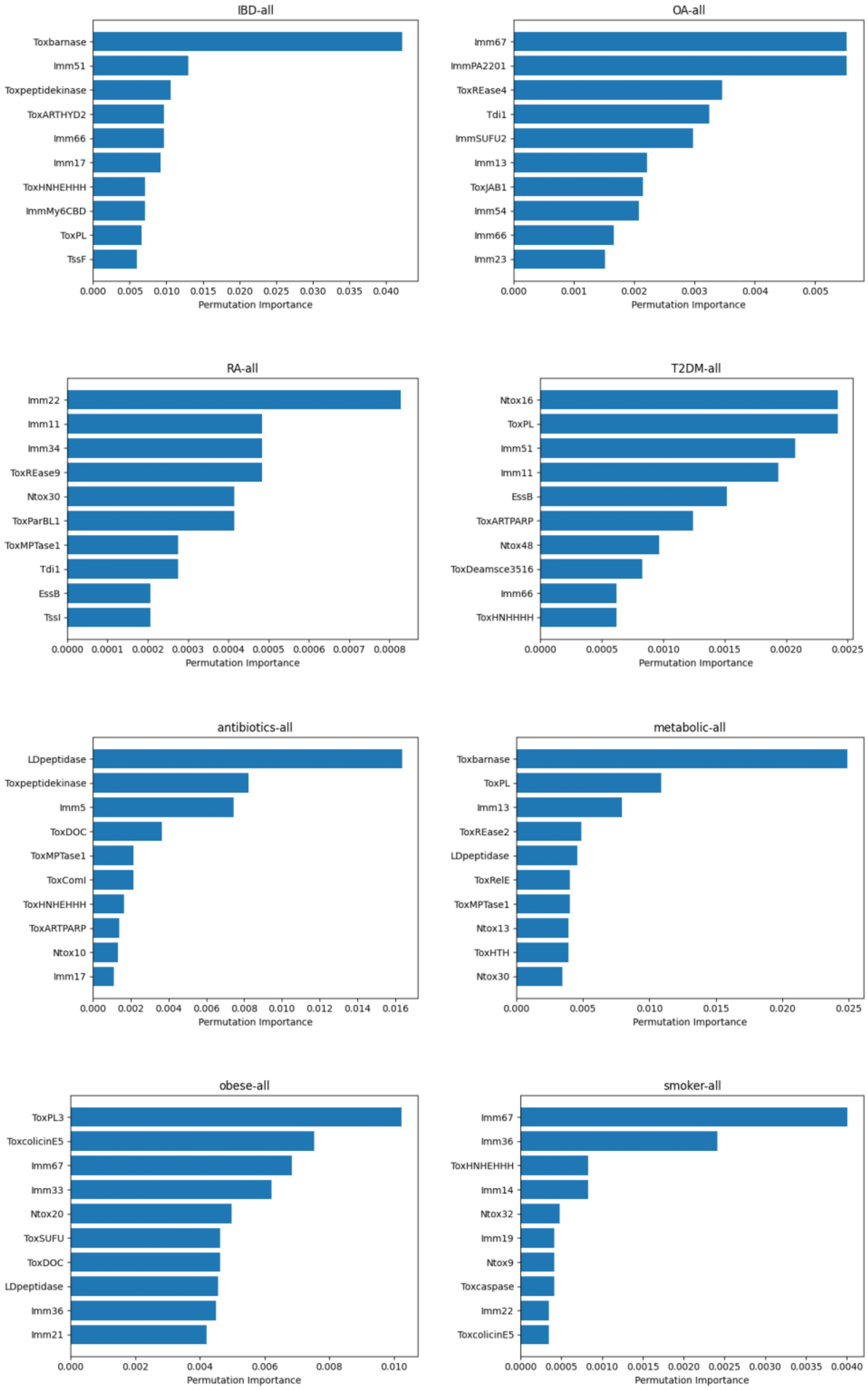


**Figure S10.** *Top distinguishing features for decision tree models of each disease vs. all other diagnoses.* Importance of features in XGBoost models was quantified with permutation-based testing. The top 10 features for each diagnosis are shown.

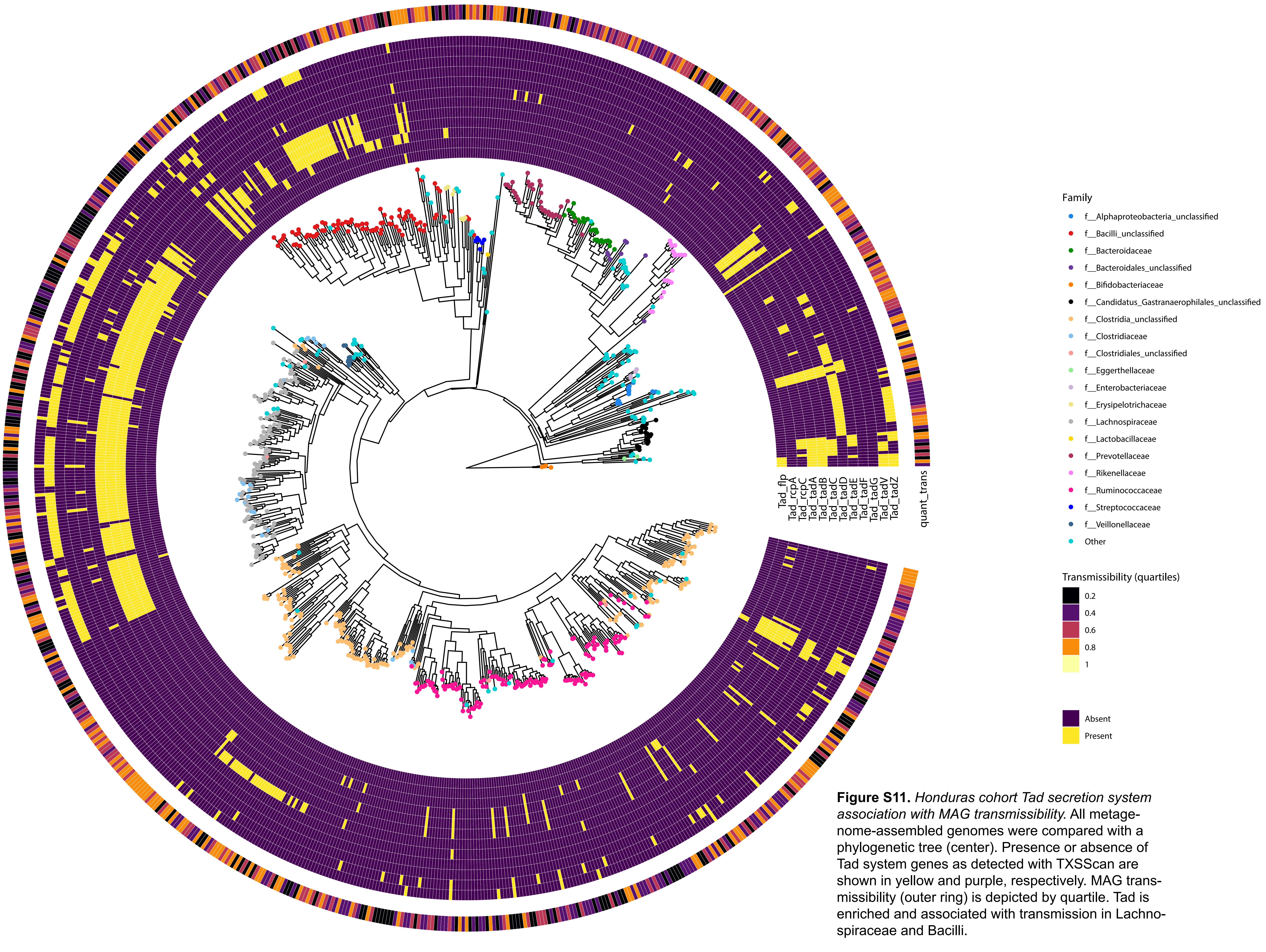


**Figure S11.** *Honduras cohort Tad secretion system association with MAG transmissibility.* All metagenome-assembled genomes were compared with a phylogenetic tree (center). Presence or absence of Tad system genes as detected with TXSScan are shown in yellow and purple, respectively. MAG transmissibility (outer ring) is depicted by quartile. Tad is enriched and associated with transmission in Lachnospiraceae and Bacilli.

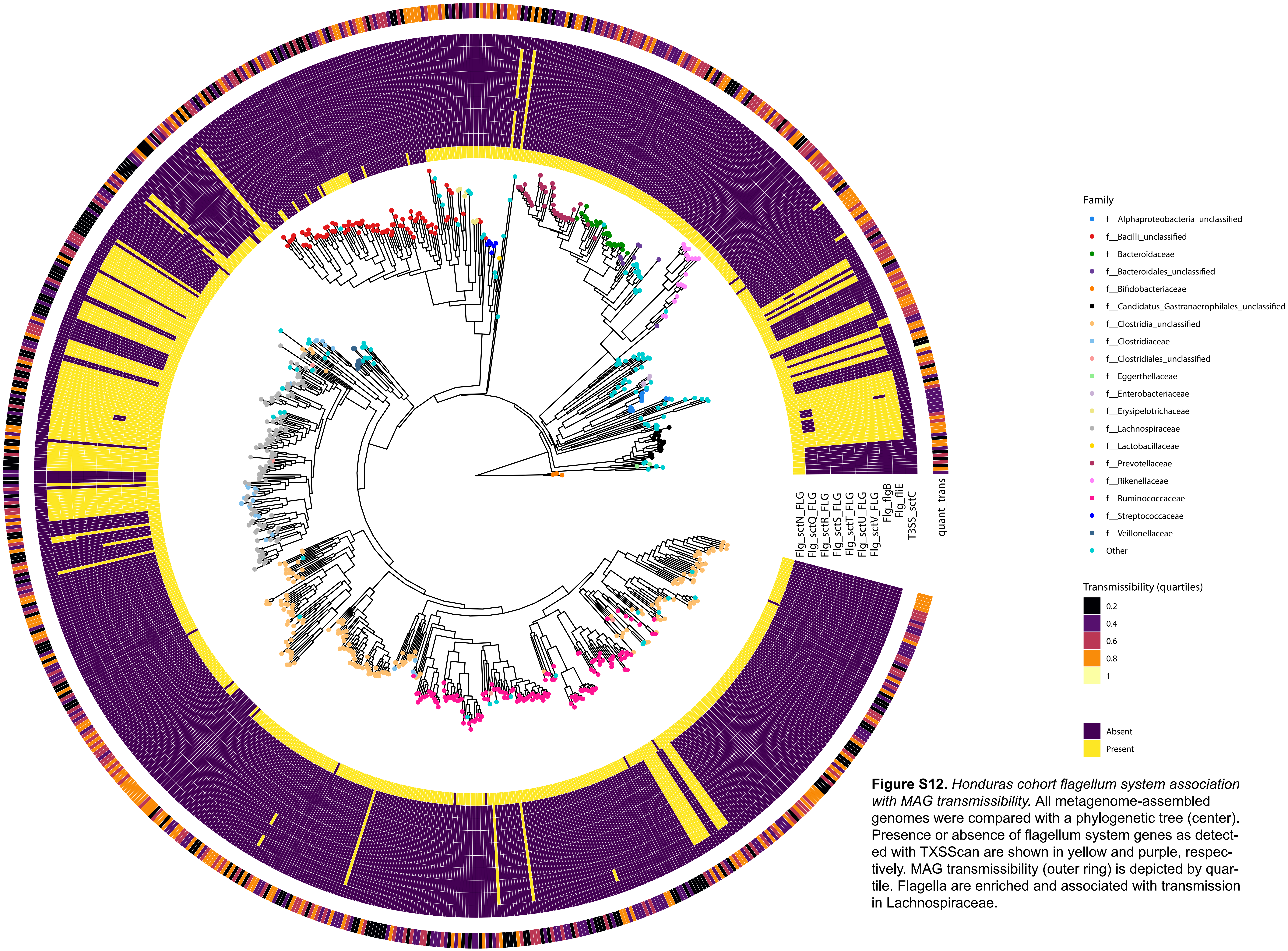


**Figure S12.** *Honduras cohort flagellum system association with MAG transmissibility.* All metagenome-assembled genomes were compared with a phylogenetic tree (center). Presence or absence of flagellum system genes as detected with TXSScan are shown in yellow and purple, respectively. MAG transmissibility (outer ring) is depicted by quartile. Flagella are enriched and associated with transmission in Lachnospiraceae.

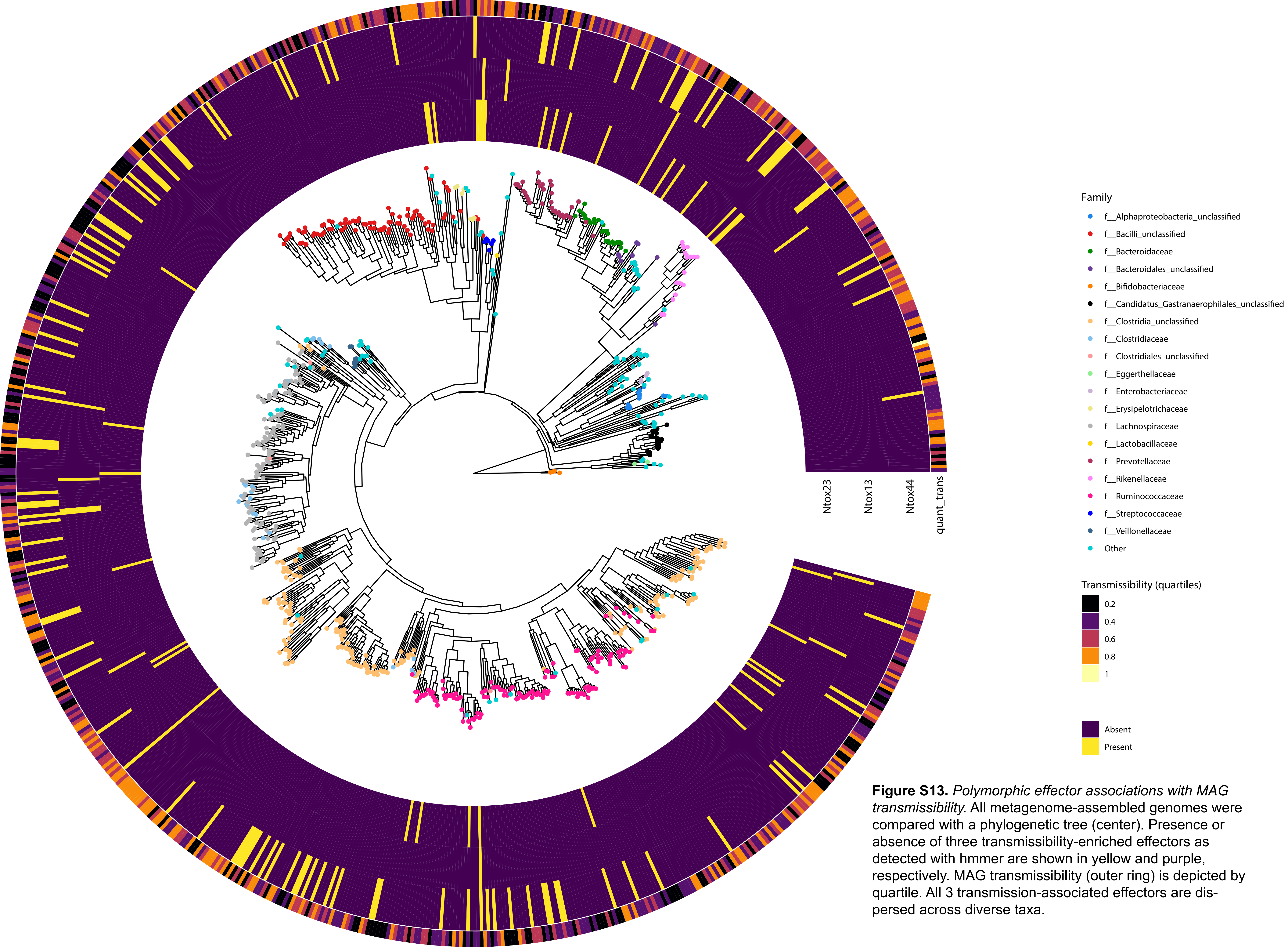


**Figure S13.** *Polymorphic effector associations with MAG transmissibility.* All metagenome-assembled genomes were compared with a phylogenetic tree (center). Presence or absence of three transmissibility-enriched effectors as detected with hmmer are shown in yellow and purple, respectively. MAG transmissibility (outer ring) is depicted by quartile. All 3 transmission-associated effectors are dispersed across diverse taxa.
